# Sequence and structural diversity of mouse Y chromosomes

**DOI:** 10.1101/096297

**Authors:** Andrew P Morgan, Fernando Pardo-Manuel de Villena

**Author notes:** Corresponding author: 5049C Genetic Medicine Building Department of Genetics University of North Carolina Chapel Hill NC 27599-7264.

## Abstract

Over the 180 million years since their origin, the sex chromosomes of mammals have evolved a gene repertoire highly specialized for function in the male germline. The mouse Y chromosome is unique among mammalian Y chromosomes characterized to date in that it is large, gene-rich and euchromatic. Yet little is known about its diversity in natural populations. Here we take advantage of published whole-genome sequencing data to survey the diversity of sequence and copy number of sex-linked genes in three subspecies of house mice. Copy number of genes on the repetitive long arm of both sex chromosomes is highly variable, but sequence diversity in non-repetitive regions is decreased relative to expectations based on autosomes. We use simulations and theory to show that this reduction in sex-linked diversity is incompatible with neutral demographic processes alone, but is consistent with recent positive selection on genes active during spermatogenesis. Our results support the hypothesis that the mouse sex chromosomes are engaged in ongoing intragenomic conflict.

## Introduction

Sex chromosomes have emerged many times in independent plant and animal lineages. The placental mammals share a sex chromosome pair that originated approximately 180 million years ago (Mya) (Hughes and Page 2015). In the vast majority of mammal species, the Y chromosome is sex-determining: presence of the Y-encoded protein SRY is sufficient to initiate the male developmental program (Berta et al. 1990). Since their divergence from the ancestral X chromosome, mammalian Y chromosomes have lost nearly all of their ancestral gene content (Figure 1A). Although these losses have occurred independently along different lineages within the mammals, the small subset of genes that are retained in each linage tend to be dosage-sensitive and have housekeeping functions in core cellular processes such as transcription and protein degradation (Bellott et al. 2014; Cortez et al. 2014). Contrary to bold predictions that the mammalian Y chromosome is bound for extinction (Graves 2006), empirical studies of Y chromosomes have demonstrated that most gene loss occurs in early proto-sex chromosomes, and that the relatively old sex chromosomes of mammals are more stable (Bellott et al. 2014). The evolutionary diversity of Y chromosomes in mammals arises from the set of Y-acquired genes, which make up a small fraction of some Y chromosomes and a much larger fraction in others — from 5% in rhesus to 45% in human (Hughes and Page 2015) (Figure 1B). These genes are often present in many copies and are highly specialized for function in the male germline (Lahn and Page 1997; Soh et al. 2014).

**Figure 1:**
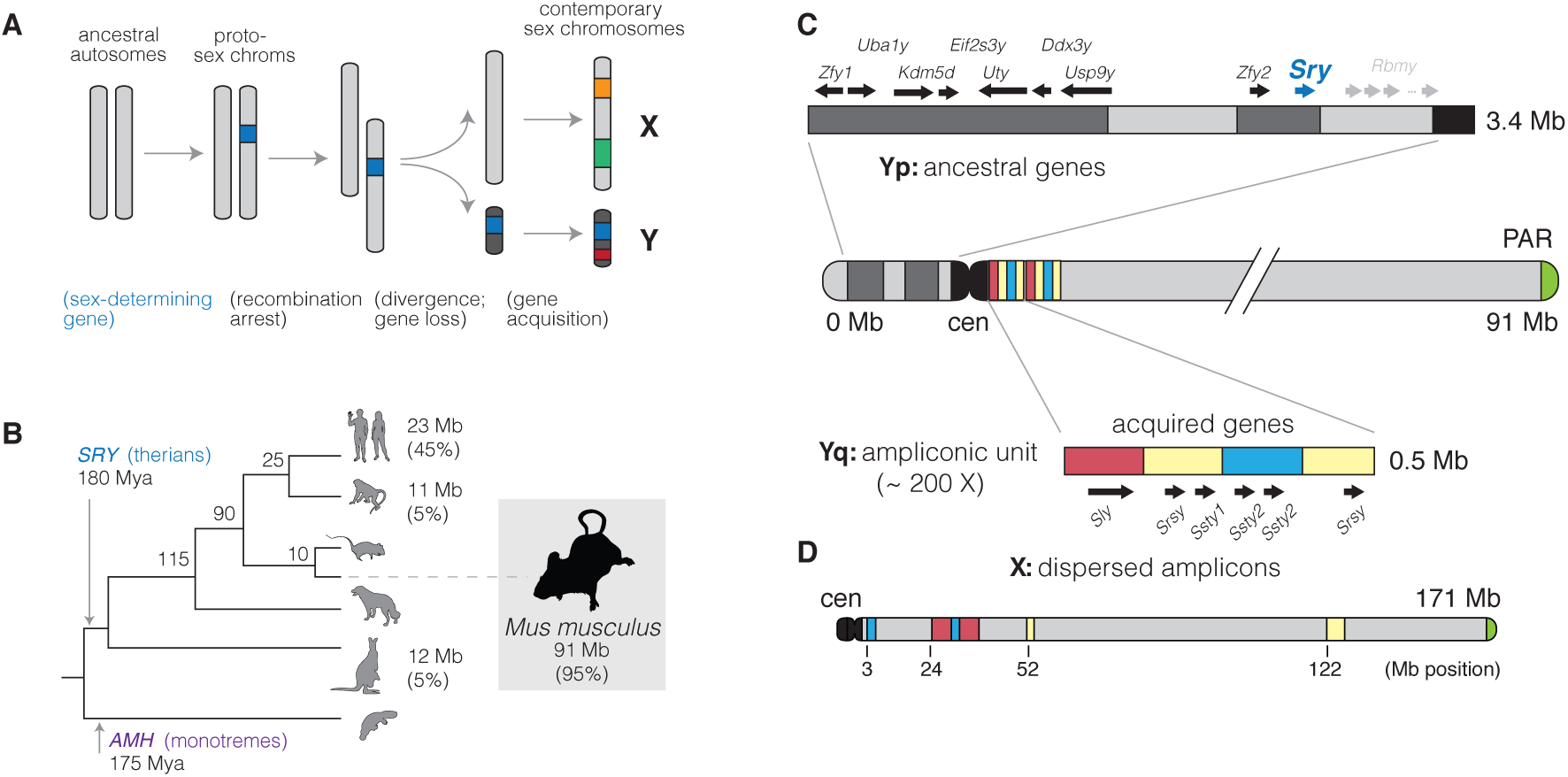
Evolution of mammalian Y chromosomes. (**A**) Evolution of heteromorphic sex chromosomes. (**B**) Y chromosomes of mammals. The Y chromosome of therian mammals, characterized by the sex-determining factor *SRY*, diverged from the X chromosome approximately 180 Mya. (The monotremata have a different sex-determining factor, *AMH*, and an idiosyncratic five-pair sex chromosome system.) Y chromosome sizes and the fraction of sequence occupied by multicopy, Y-acquired genes are shown at the tips of the tree. (**C**) Structure of the Y chromosome in the C57BL/6J reference strain. The short arm of the Y chromosome (Yq) consists primarily of genes shared with the X chromosome and retained since the sex chromosomes diverged from the ancestral autosome pair. These genes are interspersed with blocks of segmental duplications (light grey). The sex-determining factor *Sry* is encoded on the short arm. The long arm (Yq) consists of approximately 200 copies of a 500 kb repeating unit containing the acquired genes *Sly*, *Ssty1*, *Ssty2* and *Srsy*. The sequence in the repeat unit can be roughly divided into three families “red,” “yellow” and “blue” following (Soh et al. 2014). (**D**) The X choromosome, un-like the Y chromosome, is acrocentric. Homologs of the acquired genes on the Y chromosome (*Slx*, *Slxl1*, *Sstx* and *Srsx*; shown above using colored blocks as on the Y) are present in high copy number but are arranged in tandem chunks, rather than intermingled as on the Y.

The Y chromosome of the house mouse (*Mus musculus*) stands out among mammalian Y chromosomes both for its sheer size and its unusual gene repertoire. Early molecular studies of the mouse Y chromosome hinted that it consisted of mostly of repetitive sequences, with copy number in the hundreds, and that it was evolving rapidly (Nishioka and Lamothe 1986; Eicher et al. 1989). Unlike other mammalian Y chromosomes, which are dominated by large blocks of heterochromatin (Hughes and Page 2015), the mouse Y chromosome was also known to be large and almost entirely euchromatic. Spontaneous mutations in laboratory stocks allowed the mapping of male-specific tissue antigens and the sex-determining factor *Sry* to the short arm of the chromosome (Yp) (McLaren et al. 1988), while lesions on the long arm (Yq) were associated with infertility and defects in spermatogenesis (Styrna et al. 1991; Burgoyne et al. 1992; Touré et al. 2004).

Sequencing, assembly and annotation of the mouse Y chromosome in the inbred strain C57BL/6J was finally completed in 2014 after more than a decade of painstaking effort (Soh et al. 2014). Ancestral genes are restricted to Yp and are fewer in number on the mouse Y chromosome than in other studied mammals. Yq was shown to consist of approximately 200 copies of a 500 kb unit — the “huge repeat array” — containing the acquired genes *Sly*, *Ssty1*, *Ssty2* and *Srsy* (Figure 1C). *Sly* and its X-linked homologs *Slx* and *Slxl1* are found only in the genus *Mus* and have sequence similarity to the synaptonemal complex protein SYCP3 (Ellis et al. 2011). *Ssty1/2* and *Sstx* are most similar to members of the spindlin family (Oh et al. 1997) and are present in taxa at least as phylogenetically distant as rats. The coding potential of *Srsy* and *Srsx* is unclear, but they have sequence similarity to melanoma-related cancer/testis antigens typified by the human MAGEA family. Their phylogenetic origins remain unresolved. The genes of the huge repeat array are expressed almost exclusively in post-meiotic round spermatids and function in chromatin condensation and sperm maturation (Burgoyne et al. 1992; Touré et al. 2004, 2005; Yamauchi et al. 2009, 2010).

Independent amplification of homologous genes on the X and Y chromosomes is thought to be a byproduct of competition between the X and Y chromosomes for transmission to the next generation. The current consensus favors an unidentified X-linked sex-ratio distorter whose action is suppressed by one or more Y-linked factors (Ellis et al. 2011). Consistent with this hypothesis, SLY acts directly to maintain transcriptional repression of post-meiotic sex chromatin (PSCR, Hendriksen et al. (1995)) by recruiting a suite of repressive histone marks (Ellis et al. 2005; Cocquet et al. 2009; Moretti et al. 2016); its action is opposed by SLX and SLXL1. Imbalance between SLY and SLX/SLXL1 tilts the progeny sex ratio in favor of the overexpressing chromosome and causes defects in sperm morphology and sperm count (Touré et al. 2004; Cocquet et al. 2009, 2010). Disruption of PSCR and the related process of meiotic sex chromosome inactivation (MSCI) is also associated with male sterility in inter-subspecific hybrids between *M. m. domesticus* and *M. m. musculus* (Good et al. 2010; Campbell et al. 2013; Bhattacharyya et al. 2013; Larson et al. 2016a). Together these observations suggest that the intragenomic conflict between the sex chromosomes in mouse is played out in post-meiotic spermatids and may have mechanistic overlap with hybrid male sterility.

Intragenomic conflict can have a profound impact on the genetic diversity of sex chromosomes in natural populations. Sex-ratio-distorter systems in *Drosophila* provide some of the best-known examples (Jaenike 2001; Derome et al. 2004; Kingan et al. 2010). The extent to which diversity on mouse sex chromosomes is influenced by intragenomic conflict remains an open question. The differential impact of selection on mouse X chromosome versus autosomes (the “faster-X” effect) is well-studied, mostly through the lens of speciation (Torgerson and Singh 2003; Kousathanas et al. 2014; Larson et al. 2016a,b). Larson et al. (2016b) used pairwise comparisons between wild-derived strains of *M. m. musculus* and *M. m. domesticus* to show that the “faster-X” effect is most prominent in two groups of genes: those expressed primarily in the testis and early in spermatogenesis (before MSCI), and those up-regulated in spermatids (after PSCR). The former set of genes is also prone to aberrant expression in sterile hybrids Larson et al. (2016a). By contrast, selective pressures imposed by intragenomic conflict between the sex chromosomes should be exerted in spermatids after the onset of PSCR. Genes with spermatid-specific expression are expected to respond most rapidly, while those with broad expression are expected to be constrained by putative functional requirements in other tissues or cell types.

In this manuscript we take advantage of the relatively recent high-quality assembly of the mouse Y chromosome (Soh et al. 2014) and public sequencing data from a diverse sample of wild mice to perform a survey of sequence and copy-number diversity on the sex chromosomes. We use complementary gene-expression data and annotations to partition the analysis into functionally-coherent groups of loci. We find that sequence diversity is markedly reduced on both the X and Y chromosomes relative to expectations for a stationary population. This reduction cannot be fully explained by any of several demographic models fit to autosomal data, but Y-linked diversity in *M. m. domesticus* is consistent with a recent selective sweep on Y chromosomes. Copy number of genes expressed in spermatids supports the hypothesis that intragenomic conflict between the sex chromosomes during spermiogenesis is an important selective pressure. These analyses broaden our understanding of the evolution of sex chromosomes in murid rodents and support an important role for positive selection in the male germline.

## Results

### A survey of Y-linked coding variation in mouse

Whole-genome or whole-exome sequence data for 91 male mice was collected from published sources (Keane et al. 2011; Doran et al. 2016; Harr et al. 2016; Morgan et al. 2016a; Neme and Tautz 2016; Sarver et al. 2017). The final set consists of 62 wild-caught mice; 21 classical inbred strains; and 8 wild-derived inbred strains (Table 1 and **Table S1**). The three cardinal subspecies of *M. musculus* (*domesticus*, *musculus* and *castaneus*) are all represented, with *Mus spretus* and *Mus spicilegus* as close outgroups and *Mus caroli*, *Mus cookii*, and *Nannomys minutoides* as more distant outgroups. Our sample spans the native geographic range of the house mouse and its sister taxa (Figure 2A).

**Table 1:** Wild and laboratory mice used in this study.

| Type | Population | Country | Males | Females |
| --- | --- | --- | --- | --- |
| wild | *M. m. domesticus* | DE | 8 | 1 |
|  |  | FR | 7 | 0 |
|  |  | IR | 5 | 0 |
|  | *M. m. musculus* | AF | 5 | 1 |
|  |  | CZ | 2 | 5 |
|  |  | KZ | 2 | 4 |
|  | *M. m. castaneus* | IN | 3 | 7 |
|  | *M. spretus* | ES | 4 | 2 |
|  |  | MA | 1 | 0 |
|  | *M. macedonicus* | MK | 1 | 0 |
|  | *M. spicilegus* | HU | 1 | 0 |
|  | *M. caroli* | TH | 0 | 1 |
|  | *M. cookii* | TH | 1 | 0 |
|  | *Nannomys minutoides* | KE | 1 | 0 |
| wild-derived | *M. m. domesticus* | IT | 1 | 0 |
|  |  | US | 1 | 1 |
|  | *M. m. musculus* | CZ | 1 | 1 |
|  | *M. m. castaneus* | TH | 1 | 1 |
|  | *M. spretus* | ES | 1 | 0 |
| classical lab | - | - | 21 | 4 |

**Figure 2:**
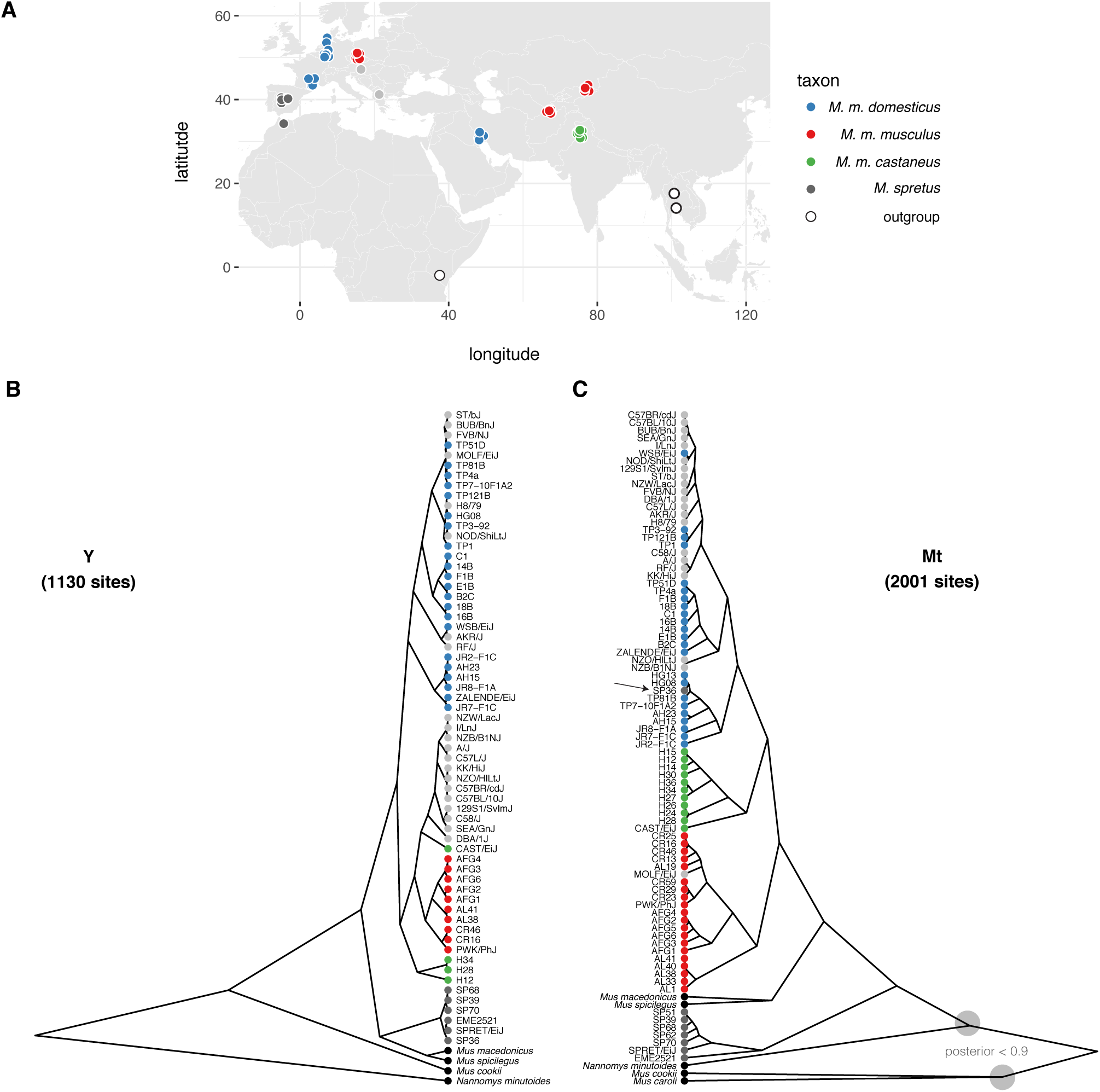
Patrilineal and matrilineal phylogeography in a geographically-diverse sample from the genus *Mus*. (**A**) Sampling locations of mice used in this study. (**B**) Phylogenetic tree from coding sites on the Y chromosome. Samples are colored according to nominal ancestry; laboratory strains are shown in light grey. (**C**) Phylogenetic tree from coding sites on the mitochondrial genome. Deep nodes with posterior support < 0.9 indicated with shaded circles.

Single-nucleotide variants (SNVs) and small indels were ascertained in 41.6 kb of sequence on Yp targeted by the Roche NimbleGen exome-capture array. To mitigate the effect of alignment errors and cryptic copynumber variation on our analyses, we discarded sites with evidence heterozygosity; fewer than 60 samples with a called genotype; or evidence of strand bias (see **Materials and methods**). In total we identified 1, 136 SNVs and 128 indels, with transition:tranversion ratio 2.1.

One group of inbred strains in our dataset — C57BL/6J (reference genome), C57BL/10J, C57L/J and C57BR/cdJ — have a known common ancestor in the year 1929, and a common ancestor with the strain C58/J in 1915 (Beck et al. 2000). Assuming an average of three generations per year, the total branch length of the pedigree connecting the C57 and C58 strains is 5, 280 generations, during which time 3 mutations occurred. We used these values to obtain a direct estimate of the male-specific point mutation rate: 1.8 × 10^−8^ (95% Poisson CI 4.5 × 10^−9^ − 4.7 × 10^−8^) bp^−1^ generation^−1^. This interval contains the sex-averaged autosomal rate of 5.4 × 10^−9^ bp^−1^ generation^−1^ recently estimated from whole-genome sequencing of mutation-accumulation lines (Uchimura et al. 2015). Using the ratio between paternal to maternal mutations in mouse estimated in classic studies from Russell and colleagues (2.78; reviewed in Drost and Lee (1995)), that estimate corresponds to male-specific autosomal rate of 7.9 × 10^−9^ bp^−1^ generation^−1^, again within our confidence interval. We note that these estimates assume that selection has been negligible in laboratory colonies.

### Phylogeny of Y chromosomes recovers geographic relationships

Phylogenetic trees for exonic regions of the Y chromosome and mitochondrial genome were constructed with BEAST (Figure 2B). The estimated time to most recent common ancestor (MRCA) of *M. musculus* Y chromosomes is 900, 000 years ago (95% highest posterior density interval [HPDI] 100, 000 − 1, 800, 000) years ago. Within *M. musculus*, the *domesticus* subspecies diverges first, although the internal branch separating it from the MRCA of *musculus* and *castaneus* is very short. Consistent with several previous studies, we find that the “old” classical inbred strains share a single Y haplogroup within *M. m. musculus*. This haplogroup is distinct from that of European and central Asian wild mice and is probably of east Asian origin (Bishop et al. 1985; Tucker et al. 1992; Nagamine et al. 1992). Strains related to “Swiss” outbred stocks (FVB/NJ, NOD/ShiLtJ, HR8) and those of less certain American origin (AKR/J, BUB/BnJ) (Beck et al. 2000) have Y chromosomes with affinity to western European populations. *M. m. castaneus* harbors two distinct and paraphyletic lineages: one corresponding to the Indian subcontinent and another represented only by the wild-derived inbred strain CAST/EiJ (from Thailand). The latter haplogroup corresponds to a southeast Asian lineage identified in previous reports that sampled more extensively from that geographic region (Geraldes et al. 2008; Yang et al. 2011). It remains unclear whether this haplogroup originated in *M. m. musculus* and displaced the *M. m. castaneus* Y chromosome in southeast Asia; or instead represents a deep branching within the (large and unsampled) population ancestral to *musculus* and *castaneus* in central Asia.

The Y-chromosome tree otherwise shows perfect concordance between clades and geographic locations. Within the *M. m. domesticus* lineage we can recognize two distinct haplogroups corresponding roughly to western Europe and Iran and the Mediterranean basin, respectively. Similarly, within *M. m. musculus*, the eastern European mice (from Bavaria, Czech Republic) are well-separated from the central Asian mice (Kazakhstan and Afghanistan). Relationships between geographic origins and phylogenetic affinity are considerably looser for the mitochondrial genome. We even found evidence for inter-specific hybridization: one nominally *M. spretus* individual from central Spain (SP36) carries a *M. spretus* Y chromosome and a *M. m. domesticus* mitochondrial genome (arrowhead in Figure 2B). Several previous studies have found evidence for introgression between *M. musculus* and *M. spretus* where their geographic ranges overlap (Orth et al. 2002; Song et al. 2011; Liu et al. 2015).

### Copy-number variation is pervasive on the Y chromosome

We examined copy number along Yp using depth of coverage. Approximately 779 kb (24%) of Yp consists of segmental duplications or gaps in the reference assembly (Figure 1); for duplicated regions we scaled the normalized read depth by the genomic copy number in the reference sequence to arrive at a final copy-number estimate for each individual. All of the known duplications on Yp are polymorphic in laboratory and natural populations (Figure 3A). The distribution of CNV alleles follows the SNV-based phylogenetic tree. Only one CNV region on Yp, adjacent to the centromere, contains a known protein-coding gene (*Rbmy*). Consistent with a previous report (Ellis et al. 2011), we find that *musculus* Y chromosomes have more copies of *Rbmy* than *domesticus* or *castaneus* chromosomes.

**Figure 3:**
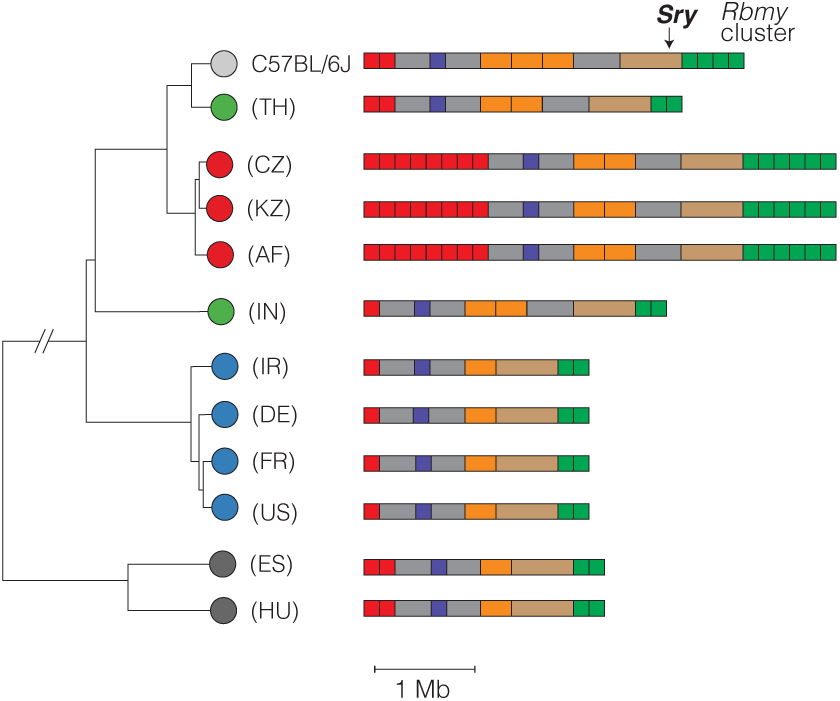
Copy-number variation on Yp. Schematic view of copy-number variable regions of the Y chromosome short arm (Yp) superposed on SNV-based phylogenetic tree. All CNVs shown overlap a segmental duplication in the reference sequence (strain C57BL/6J). One CNV overlaps a known protein-coding gene: an expansion of the ampliconic *Rbmy* cluster (green) in *M. m. musculus*. Color scheme for *Mus* taxa follows **Figure 2**.

The highly repetitive content of Yq precludes a similarly detailed characterization of copy-number variation along this chromosome arm. However, we can estimate the copy number of each of the three gene families present (*Sly*, *Ssty1/2* and *Srsy*) by counting the total number of reads mapped to each and normalizing for sequencing depth. The hypothesis of X-Y intragenomic conflict predicts that, if expression levels are at least roughly proportional to copy number, amplification of gene families on Yq should be countered by amplification of their antagonistic homologs on the X chromosome (or vice versa.) We tested this hypothesis by comparing the copy number of X- and Y-linked homologs of the *Slx/y*, *Sstx/y* and *Srsx/y* families in wild mice. Figure 4 shows that copy number on X and Y chromosomes are indeed correlated for *Slx/y*. The relationship between *Slx*-family and *Sly*-family copy number is almost exactly linear (slope = 0.98 [95% CI 0.87 − 1.09]; *R*^2^ = 0.87). We note that samples are not phylogenetically independent, so the statistical significance of the regression is exaggerated, but the qualitative result clearly supports previous evidence that conflict between X and Y chromosomes is mediated primarily through *Slx* and *Sly* (Cocquet et al. 2012). Size differences estimated from *Sly* copy number are also concordant with cytological observations that the Y chromosomes of wild-caught *M. m. musculus* appear much larger than those of *M. spicilegus* or *M. spretus* (Bulatova and Kotenkova 1990; Yakimenko et al. 1990).

**Figure 4:**
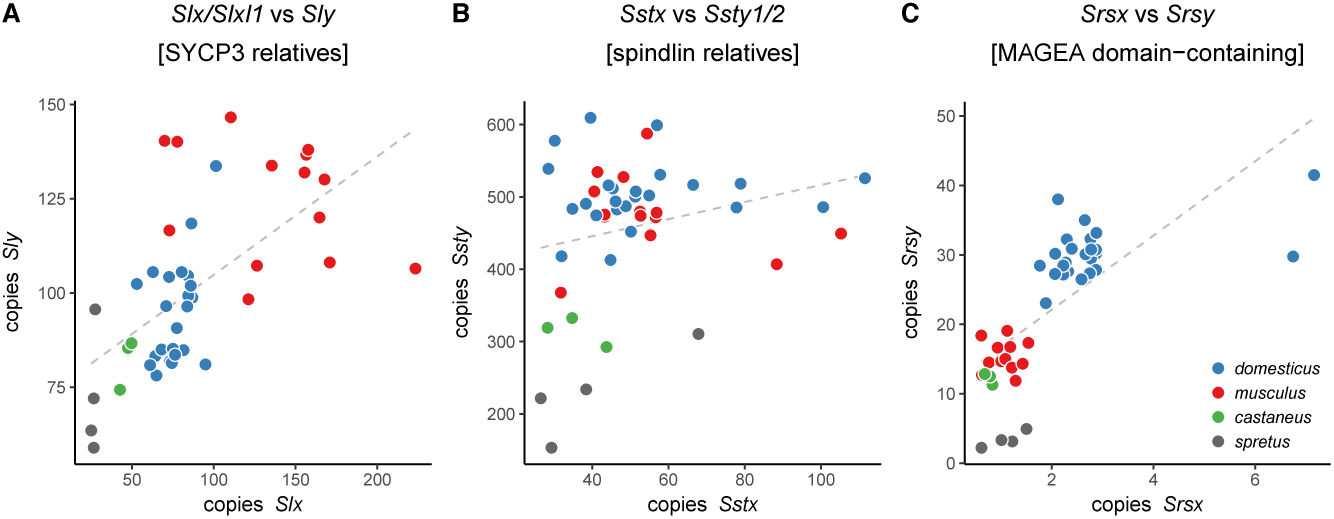
Approximate copy number of co-amplified gene families on X and Yq. Each dot represents a single individual. Grey dashed line is simple linear regression of Y-linked versus X-linked copy number.

It has recently been shown that two regions of the autosomes — on chromosomes 5 and 14 — have a suite of epigenetic marks similar to the sex chromosomes in post-meiotic spermatids (Moretti et al. 2016). These autosomal regions harbor many copies of a family of genes (known alternatively as *Speer* (Spiess et al. 2003) or *α*-takusan (Tu et al. 2007)) expressed in spermatids. The copy number of *Speer* family members is, like *Sly*, correlated with that of *Slx/Slxl1* (Figure S1). This finding supports the hypothesis that the *Speer* family may be involved in sex-chromosome conflict in spermatids.

The scale of copy number change within the *M. musculus* lineage suggests a high underlying mutation rate. We used whole-genome sequence data from a panel of 69 recombinant inbred lines (RILs) from the Collaborative Cross (CC; Srivastava et al. (2017)) to estimate the rate of copy-number change on Yq. Each CC line is independently derived from eight inbred founder strains via two generations of outcrossing followed by sibling matings until inbreeding is achieved (Consortium 2012). Distinct CC lines inheriting a Y chromosome from the same founder strain thus share an obligate male ancestor in the recent past, but no more recently than the start of inbreeding (Figure 5A). We estimated read depth in 100 kb bins across Yq and normalized each bin against the median for CC lines inheriting a Y chromosome from the same founder strain. This normalization effectively removes noise from mapping of short reads to repetitive sequence and uncovers CNVs from 6 to 30 Mb in size in 5 CC lines carrying 3 different Y chromosomes (Table 2, **Table S2** and Figure 5B). Because the pedigree of each CC line is known, mutation rates — for each Y haplogroup, and overall — can be estimated directly, assuming each new allele corresponds to a single mutational event. Our estimate of 0.30 (95% Poisson CI 0.098 − 0.70) mutations per 100 father-son transmissions is about tenfold higher than ampliconic regions of the human Y chromosome (Repping et al. 2006), and places the mouse Yq among the most labile sequences known in mouse or human (Egan et al. 2007; Itsara et al. 2010; Morgan et al. 2016b). New Yq alleles also provide opportunities to investigate the effects of Yq copy number on fertility, sperm phenotypes and sex ratio (as in, among others, Styrna et al. (1991); Touré et al. (2004); Yamauchi et al. (2010); Cocquet et al. (2012); Fischer et al. (2016)).

**Table 2:**
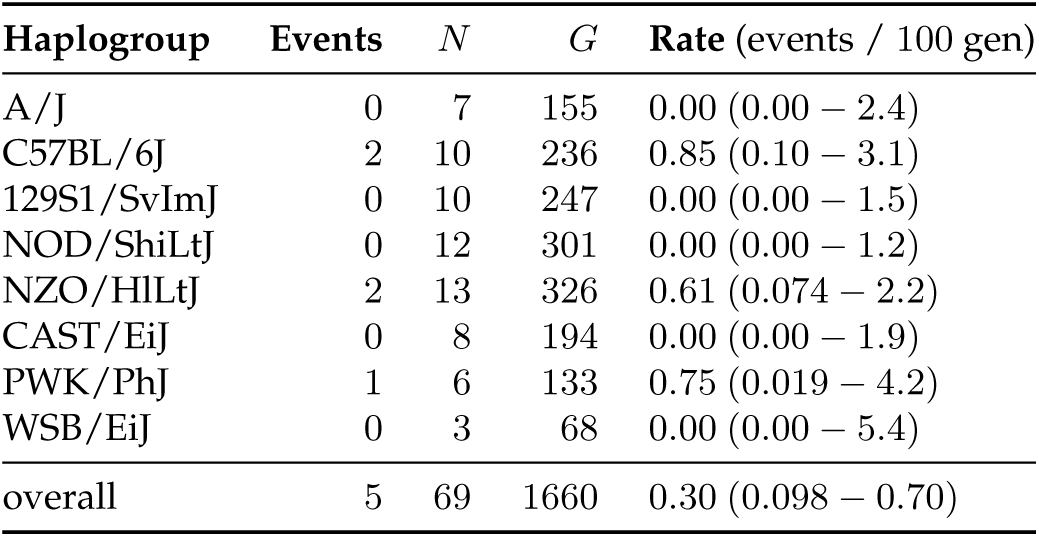
Pedigree-based estimates of mutation rates on Yq. *N*, number of CC lines with each Y chromosome haplogroup; *G*, total number of breeding generations.

**Figure 5:**
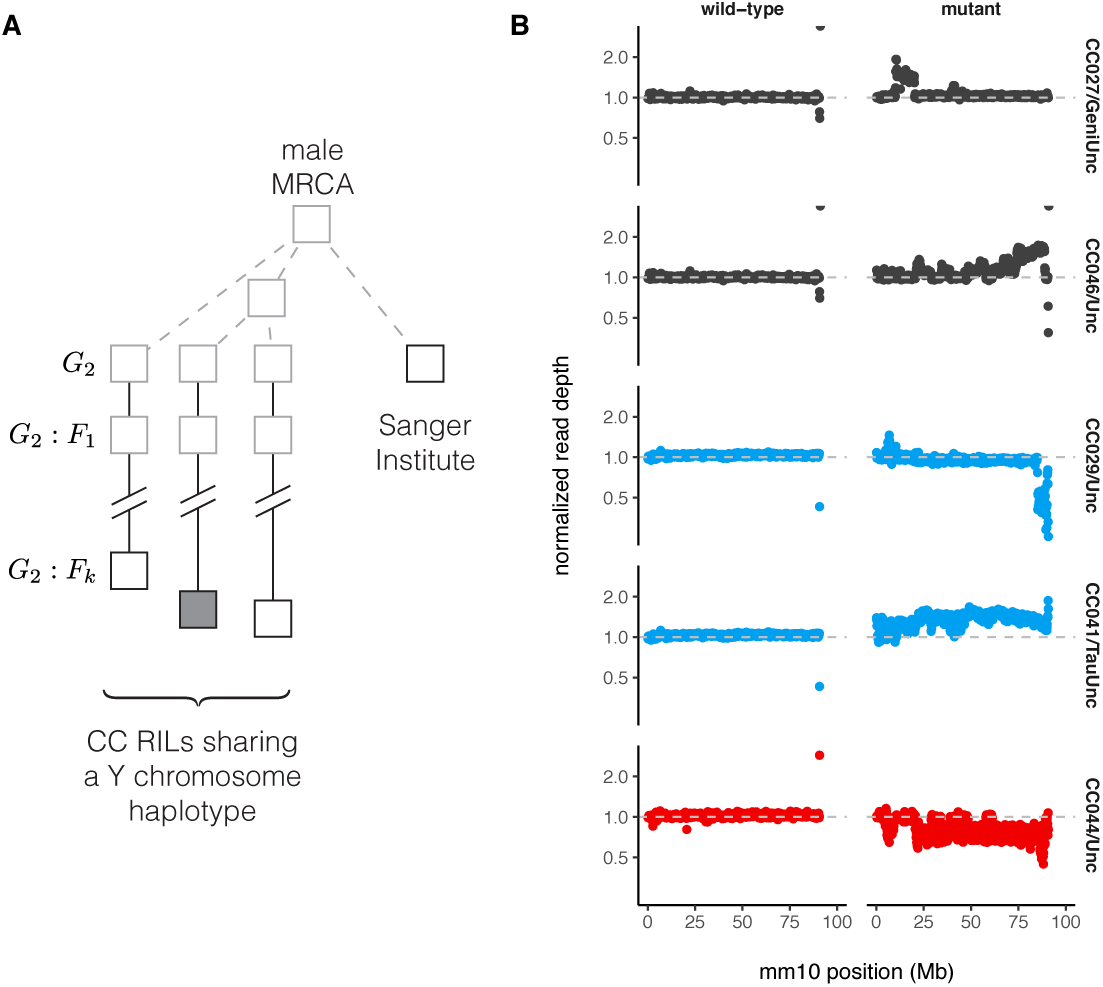
Copy-number variation on Yq in the Collaborative Cross. (**A**) Pedigree-based estimates of mutation rate on the Y chromosome long arm (Yq). Multiple recombinant inbred lines (RILs) from the Collaborative Cross (CC) panel share the same Y chromosome haplotype, with (filled shape) or without (open shape) a putative *de novo* CNV. These Y chromosome lineages are separated from their common male ancestor by an unknown number of generations prior to the initiation of the CC (grey dashed lines), plus a known number of generations of CC breeding (solid lines.) Representatives of the founder strains of the CC were sequenced at the Sanger institute; the number of generations separating the Sanger mouse from the common male ancestor is also unknown. (**B**) Normalized read depth across Yq for CC lines with *de novo* CNVs on Yq. Points are colored according to founder Y chromosome haplogroup. A representative wild-type line is shown for each mutant.

### Sequence diversity is markedly reduced on both sex chromosomes

We next used whole-genome sequence data to examine patterns of nucleotide diversity within *Mus musculus* in non-repetitive sequence on Yp compared to the autosomes and X chromosome. To do so we first identified a subset of wild mice without evidence of cryptic relatedness (see **Materials and methods**); this left 20 male and 1 female *M. m. domesticus* (hereafter *dom*), 9 male and 10 female *M. m. musculus* (*mus*) and 3 male and 7 female *M. m. castaneus* (*cas*). Analyses of autosomes used both males and females from each population; sex chromosome analyses used males only to avoid introducing technical artifacts associated with differences in sample ploidy. Diversity statistics were calculated from the joint site frequency spectrum (SFS), which in turn was estimated directly from genotype likelihoods rather than hard genotype calls (Korneliussen et al. 2014).

We estimated nucleotide diversity in four classes of sites: intergenic sites (putatively neutral); introns; 4-fold degenerate sites; and 0-fold degenerate sites. Putatively neutral sites are useful for estimating demographic parameters, while the latter three classes are useful for assessing the impact of selection. Sites on the sex chromosomes are subject to different selective pressures than autosomal sites, both because they are “exposed” in the hemizygous state in males and because, in mammals, the sex chromosomes are enriched for genes with sex-specific expression patterns. To evaluate these effects we further subdivided genic sites according to geneexpression patterns inferred from two expression datasets, one in eighteen adult tissues and one a time course across spermatogenesis (see **Materials and methods**). Genes on the autosomes and X chromosome were classified along two independent axes: testis-specific versus ubiquitously-expressed; or expressed early in meiosis, prior to MSCI, versus expressed in post-meiotic spermatids. (Y chromosome genes are not subdivided, since they are few in number and inherited as a single linkage block.) All diversity estimates are shown in **Table S3**. For putatively neutral sites on the autosomes, our estimates of pairwise diversity (*π*_dom_ = 0.339%, *π*_mus_ = 0.325%, *π*_cas_ = 0.875%) are consistent with previous reports based on overlapping samples (Geraldes et al. 2008; Halligan et al. 2013; Kousathanas et al. 2014; Harr et al. 2016). Within each chromosome type, levels of diversity follow the expected rank order: intergenic sites > introns ≈ 4-fold degenerate (synonymous) sites > 0-fold degenerate (non-synonymous) sites.

For the X chromosome, we further examined the relationship between sequence diversity and local sequence features including recombination rate, X-Y gametologous amplicons, gene sets described above and blocks of conserved synteny with rat (Figure S2). Diversity is reduced across the entire X chromosome in all three populations, in marked contrast to local “troughs” observed in great apes (Nam et al. 2015). Regression of pairwise diversity (*θ*_*π*_) on distance away from ubiquitously-expressed genes, meiosis genes, spermatid genes, and X-Y ampliconic genes was significant only in *musculus* for ubiquitously expressed genes (*t* = 6.6, Bonferoni-corrected *p* = 6.8 × 10^−11^). Similarly — and surprisingly — there was no relationship (*t* = −1.2, *p* = 0.23) between sequence diversity and recombination rate at 100 kb resolution, as estimated from the Diversity Outbred mouse stock (Morgan et al. 2017). (We speculate that characterizing recombination at finer scale from linkage disequilibrium (Auton and McVean 2007) would provide a more powerful test.)

In a panmictic population with equal effective number of breeding males and breeding females (*ie.* with equal variance in reproductive success between sexes), there are 3 X chromosomes and a single Y chromosome for every 4 autosomes. The expected ratios of X:A and Y:A diversity are therefore 3/4 and 1/4, respectively, if mutation rates in males and females are equal (Charlesworth et al. 1987). We estimated X:A and Y:A for putatively neutral sites and find that diversity on both sex chromosomes is markedly reduced relative to expectations in all three populations (Table 3). The effect is strongest in *M. m. domesticus* (X:A = 0.244, Y:A = 0.0858) and weakest in *M. m. musculus* (X:A = 0.563, Y:A = 0.216). The mutation rate is higher in the male than the female germline in most mammals (recently reviewed in Scally (2016)), including mice, which might contribute to differences in observed diversity between chromosomes. We used divergence between mouse and rat at synonymous, one-to-one orthologous sites (*d*_rat_) on autosomes, X and Y chromosome as a proxy for the long-term average mutation rate, and corrected X:A and Y:A estimates for differences in mutation rate (“corrected” rows in Table 3). Even with this correction, X- and Y-linked diversity remains below expectations. Scaled diversity estimates for each class of sites are shown in Figure 6. Reduction in X:A diversity has been described previously on the basis of targeted sequencing of a few loci in all thee subspecies (Baines and Harr 2007), and for *M. m. castaneus* on the basis of whole-genome sequencing (Halligan et al. 2013; Kousathanas et al. 2014). A reduction in Y:A has not, to our knowledge, been reported.

**Table 3:** Diversity ratios between pairs of chromosome types relative to neutral expectations, with 95% confidence intervals. Both raw diversity and diversity corrected for divergence to rat are shown.

| Comparison | Expected | Scaling | Population |  |  |
| --- | --- | --- | --- | --- | --- |
|  |  |  | *dom* | *mus* | *cas* |
| X:A | $3/4$ | raw | 0.244 (0.219 – 0.268) | 0.563 (0.506 – 0.613) | 0.501 (0.455 – 0.538) |
|  |  | corrected | 0.291 (0.261 – 0.319) | 0.670 (0.603 – 0.729) | 0.597 (0.542 – 0.641) |
| Y:A | $1/4$ | raw | 0.0858 (0.0805 – 0.0911) | 0.216 (0.207 – 0.227) | 0.128 (0.122 – 0.134) |
|  |  | corrected | 0.0924 (0.0867 – 0.0981) | 0.233 (0.223 – 0.244) | 0.137 (0.131 – 0.144) |

**Table 4:** Parameter estimates for models shown in Figure 7. Population sizes are given in thousands and times in units of *N*_0_; bootstrap standard errors in parentheses.

| Population | Model | Parameter |  |  |  |
| --- | --- | --- | --- | --- | --- |
| | | $N_0$ | $N_e$ | $f$ | $\tau$ |
| dom | neutral | 162 (2) | - | - | - |
|  | growth | 160 (50) | 230 (80) | - | 1.53 (0.06) |
|  | step-change | 230 (50) | 400 (700) | 2 (4) | 7 (6) |
|  | bottleneck | 164 (3) | 30 (50) | 0.2 (0.3) | 0 (2) |
| mus | neutral | 165 (2) | - | - | - |
|  | growth | 150 (40) | 500 (200) | - | 7 (5) |
|  | step-change | 150 (50) | 270 (100) | 1.8 (0.5) | 1 (3) |
|  | bottleneck | 164 (4) | 20 (8) | 0.12 (0.05) | 1 (1) |
| cas | neutral | 429 (3) | - | - | - |
|  | growth | 350 (100) | 2000 (1000) | - | 0 (2) |
|  | step-change | 300 (100) | 800 (700) | 3 (1) | 1 (4) |
|  | bottleneck | 431 (7) | 30 (10) | 0.08 (0.03) | 0.6 (0.9) |

**Figure 6:**
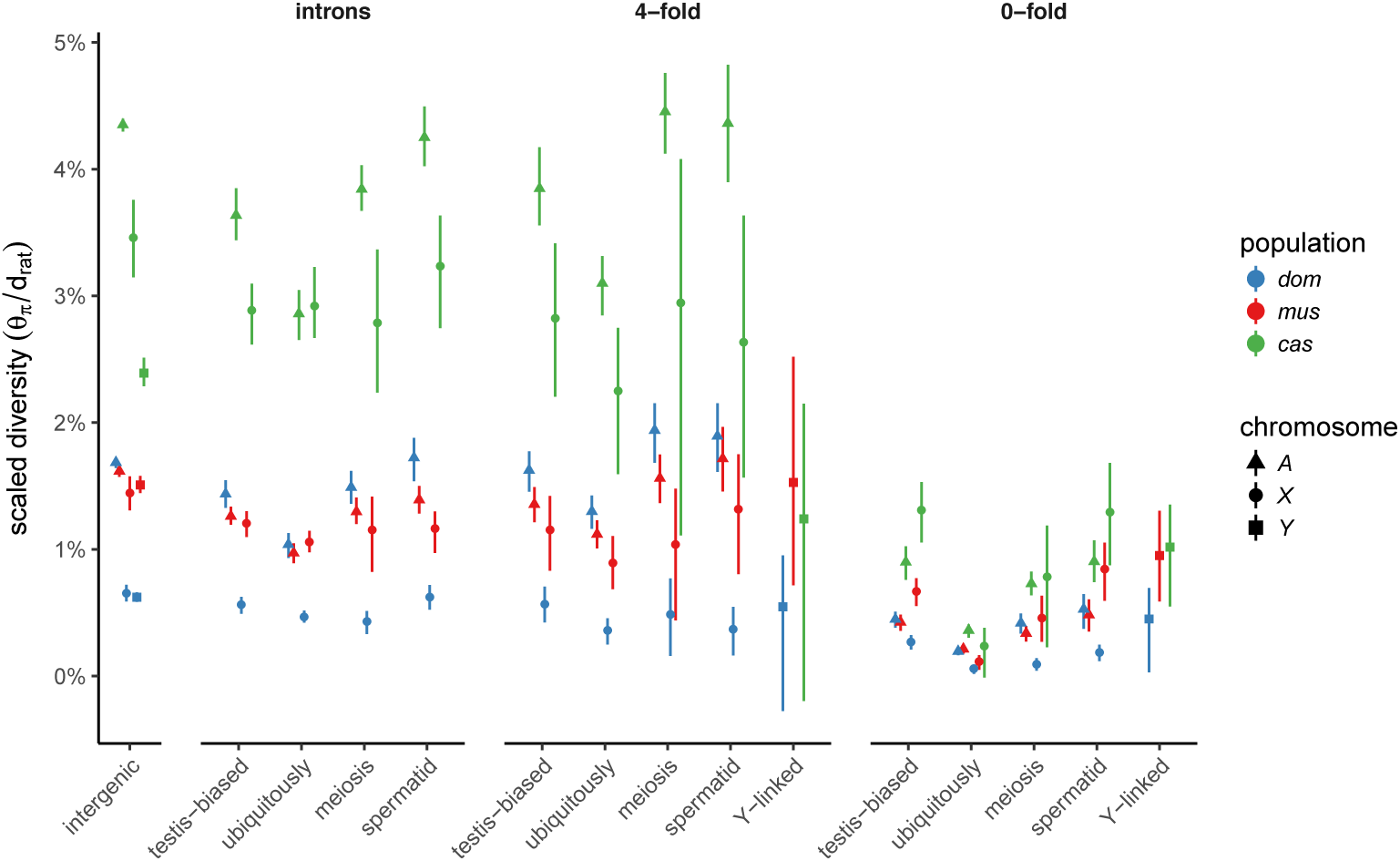
Scaled nucleotide diversity by population, site class and chromosome type. First panel from left shows estimates from intergenic sequence; remaining panels are site classes within protein-coding gene boundaries.

### Reduction in sex-linked diversity is inconsistent with simple demographic models

Sex chromosomes are affected differently than autosomes by both neutral forces, such as changes in population size (Pool and Nielsen 2007), and by natural selection (reviewed in *eg.* Ellegren (2011)). The X chromosomes of humans (Arbiza et al. 2014) and several other primate species (Nam et al. 2015) are substantially less diverse than the demographic histories of these species would predict, as a result of both purifying selection and recurrent selective sweeps. For humans, the pattern extends to the Y chromosome (Sayres et al. 2014). Having observed a deficit of polymorphism on both sex chromosomes in mouse, the central question arising in this paper is: to what extent is sex-chromosome diversity reduced by natural selection? A rich body of literature already exists for the influence of selection on the mouse X chromosome, especially in the context of speciation (Good et al. 2008; Teeter et al. 2008; Baines and Harr 2007; Kousathanas et al. 2014; Larson et al. 2016a,b), so we directed our focus to the lesser-studied Y chromosome.

To establish an appropriate null against which to test hypotheses about natural selection on the sex chromosomes, we followed an approach similar to Sayres et al. (2014). We fit four simple demographic models to SFS from putatively-neutral intergenic sites on the autosomes using the maximum-likelihood framework implemented in *∂a∂i* (Gutenkunst et al. 2009) (Figure 7A). Each model is parameterized by an initial effective population size (*N*_0_), a size change (expressed as fraction *f* of starting size for models involving instantaneous size changes, or the ending population size *N*_*e*_ for the exponential growth models), and a time of onset of size change (*τ*). The relative fit of each model was quantified using the method of Aikake weights (Akaike 1978). All four models can be viewed as nested in the family of three-epoch, piecewise-exponential histories. In principle, such models are identifiable with sample size of 4 × 3 = 12 or more chromosomes (6 diploid individuals) (Bhaskar and Song 2014). In practice, more than one model fits each population about equally well (or equally poorly), with the exception of *M. m. castaneus*, which is best described by the “step-change” model (Figure 7B-C). Of course the true history of each population is almost certainly more complex than any of our models. Our goal is not to obtain a comprehensive description of mouse population history as such, but rather to pick an appropriate null model against which we can test hypotheses about selection. For *domesticus*, the stationary model is the most parsimonious; for *musculus*, the exponential-growth model.

**Figure 7:**
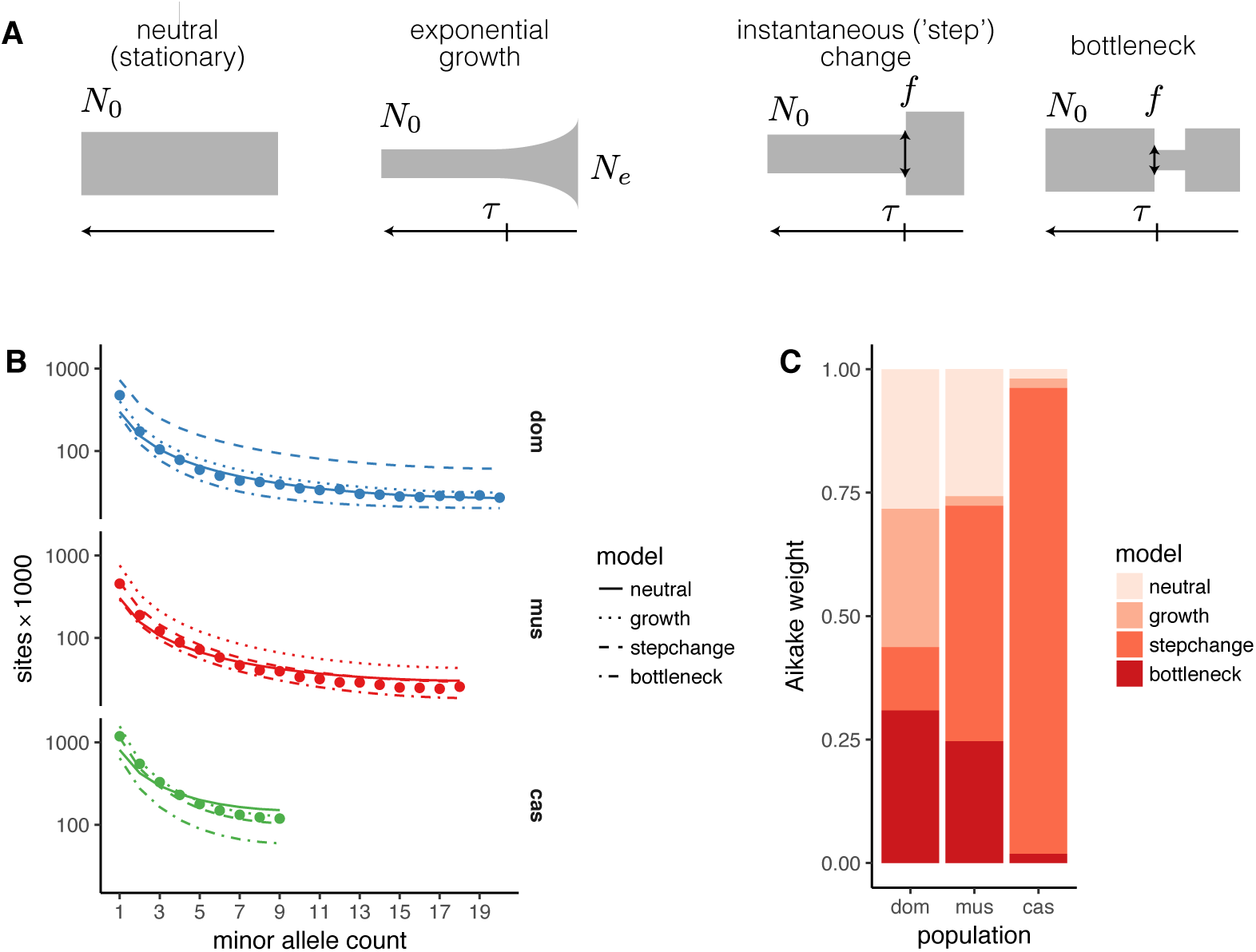
Inference of demographic histories from autosomal sites. (**A**) Four simple demographic models fit with *∂a∂i*. Each model is parameterized by one or more of an ancestral effective population size (*N*_0_), time of population-size change (*τ*), change in population size as fraction of initial size (*f*), and present effective population size (*N*_*e*_). (**B**) Observed site frequency spectra by population, with fitted spectra from the four models in panel A. (**C**) Relative support for each model, quantified by Aikake weight, by population.

We also compared our estimates of sex chromosome diversity to predictions from coalescent theory (Pool and Nielsen 2007; Polanski et al. 2017) for the four models considered above. For this analysis we focused on the ratios X:A and Y:A, which are independent of autosomal effective population size. Results are shown in Figure S3, with observed X:A and Y:A ratios superposed. We summarize some relevant trends here and refer to previous reviews (Pool and Nielsen 2007; Webster and Wilson Sayres 2016) for further details. Qualitatively, both X:A and Y:A are reduced after an instantaneous contraction in population size, eventually recovering to their stationary values after about 4*N*_*e*_ generations. For a bottleneck — a contraction followed by instantaneous recovery to the initial size — X:A and Y:A are at first sharply reduced and then increased relative to a stationary population, again returning to stationary values after about 4*N*_*e*_ generations. With exponential growth, X:A and Y:A are actually increased relative to their stationary values. These patterns are modulated by the breeding sex ratio; X:A increases and Y:A decreases when females outnumber males, and vice versa. In brief, some combination of a male-biased breeding ratio and a very strong (*f* ≪ 0.1) population contraction would be required to explain the observed reductions in X:A and Y:A in *domesticus*, with somewhat milder effects required to explain the reduction in *musculus* or *castaneus*. These histories are not consistent with population histories inferred from autosomal SFS. We hypothesize that this discrepancy is explained, at least in part, by selection.

### Both sex chromosomes have been shaped by positive selection in the male germline

We used two approaches to investigate the role of selection on the sex chromosomes. First, we used a variant of the McDonald-Kreitman test (McDonald and Kreitman 1991) to obtain a non-parametric estimate of the proportion *α* of sites fixed by positive selection (loosely, the “evolutionary rate”) in genes with different expression and inheritance patterns (Smith and Eyre-Walker 2002). The rate of adaptive evolution should be faster on the X chromosome when new mutations tend to have greater fitness effect in males than in females, to be on average recessive, or both (Charlesworth et al. 1987). We might expect genes with testis-biased expression or genes expressed during spermatogenesis to be targets of male-specific selection. Consistent with previous work on the “faster-X” effect in mouse (Kousathanas et al. 2014; Larson et al. 2016a,b), we find that a greater proportion of X-linked than autosomal substitutions are adaptive. The pattern holds in all three populations (Figure 8). In *domesticus* and *musculus*, X-linked genes whose expression is biased towards early meiosis or round spermatids evolve faster than X-linked genes with ubiquitous expression or expression across spermatogenesis. By contrast, non-ampliconic Y-linked genes — all expressed during male meiosis — have evolutionary rates closer to autosomal genes, with heterogeneity across populations. Unfortunately we cannot assess the rate of sequence evolution in ampliconic gene families on the Y chromosome using short-read data.

**Figure 8:**
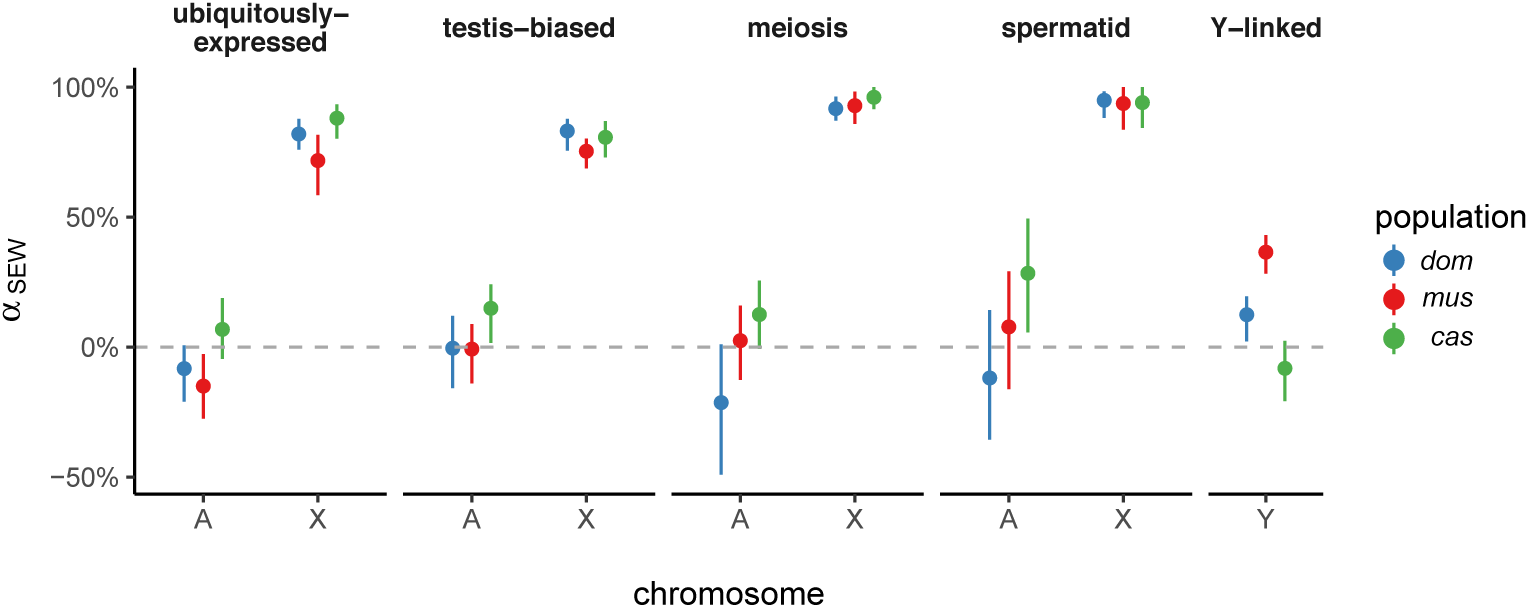
Proportion of sites fixed by positive selection (using the non-parametric estimator of Smith and Eyre-Walker (2002), *α*_SEW_) according to gene-expression class, chromosome and population. Error bars represent 95% bootstrap CIs. Ampliconic genes on X and Y are excluded.

Second, we used forward simulations from the models fit to autosomal SFS to explore the possible contribution of natural selection to the SFS of Y chromosomes. We simulated two modes of selection independently: purifying selection on linked deleterious alleles (background selection, BGS; (Hudson and Kaplan 1995)), and hard selective sweeps on newly-arising beneficial alleles. For the BGS model, we varied the proportion of sites under selection *α* and the mean population-scaled selection coefficient 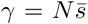; for the sweep model, we varied only the *γ* for the sweeping allele. (Simulation details are provided in **Materials and methods**.) Posterior distributions for these parameters were inferred using an approximate Bayesian computation (ABC) approach (Pritchard et al. 1999; Beaumont et al. 2002); Bayes factors were used for model comparison. The *castaneus* population was excluded from these analyses because sample size (only 3 chromosomes) was not sufficient for calculating some of the summary statistics chosen for ABC.

Results of the ABC procedure are shown in Figure S4. The Y chromosomes of *domesticus* are best approximated by the selective-sweep model. For *musculus* the result is less clear: the neutral null model actually provides the best fit, and among models with selection, the BGS model is superior. However, over the parameter ranges used in our simulations, we have limited power to discriminate between different models at the current sample size (*n* ≤ 20 chromosomes) (Figure S4B). In the best case — the selective-sweep model — we achieve only 49% recall. This reflects both the constraints of a small sample and the more fundamental limits on model identifiability for a single non-recombining locus like the Y chromosome.

If a selective sweep did occur on *domesticus* Y chromosomes, it was moderately strong: we estimate *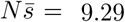* (50% HPDI 0 − 9.88) (Table 5). For comparison, *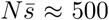* for adaptive alleles in the human lactase gene (*LCT*), a well-characterized example of recent positive selection (Tishkoff et al. 2007). Posterior distributions of several estimators of nucleotide diversity recapitulate the values observed in real data (Figure S4D). We note that, because the Y chromosome is inherited without recombination, our estimate of *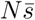* reflects the cumulative selection intensity on the entire chromosome and not necessarily on a single site.

**Table 5:**
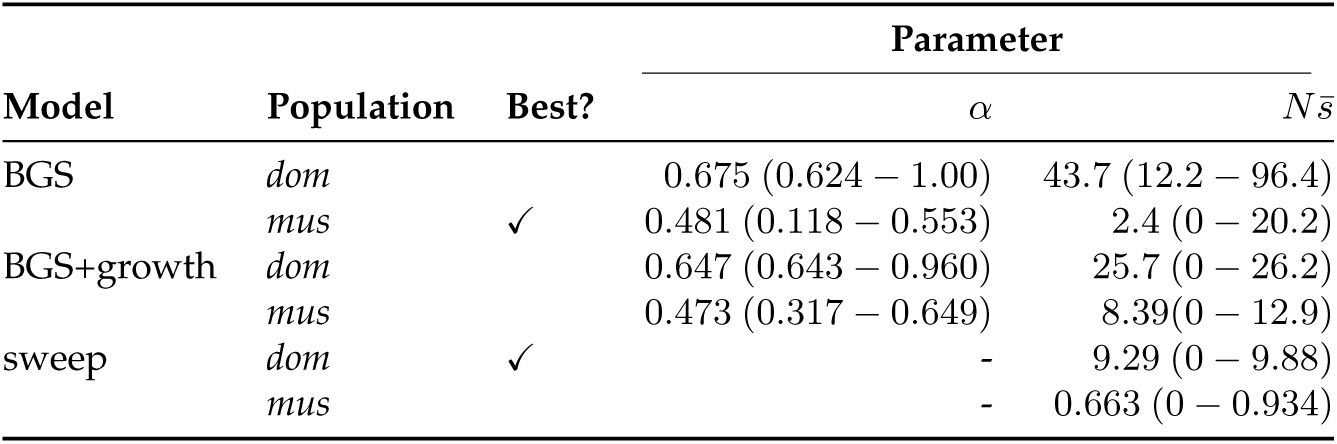
Parameter estimates from ABC. Values are shown as posterior median and 50% highest posterior density interval (HPDI). Best-fitting model for each population indicated by check mark.

**Sex-linked gene expression diverges rapidly in the testis**

Given the dramatic differences in Y-linked gene content between even closely-related *Mus* taxa, we finally asked whether patterns of gene expression showed similar divergence. In particular, we sought to test the prediction that expression patterns of Y-linked genes diverge more rapidly than autosomal genes in the testis. To that end we re-analyzed published gene expression data from the brain, liver and testis of wild-derived outbred individuals representing seven (sub)species spanning an 8 million year evolutionary transect across the murid rodents (Neme and Tautz 2016) (Figure 9A). For genes on the autosomes and X chromosome, the great majority of expression variance lies between tissues rather than between (sub)species (PC1 and PC2, cumulative 77.1% of variance explained; Figure 9B). For Y-linked genes, highly enriched for function in the male germline, most variance (PC1, 59.6% of variance explained) naturally lies between the testis and the non-germline tissues.

**Figure 9:**
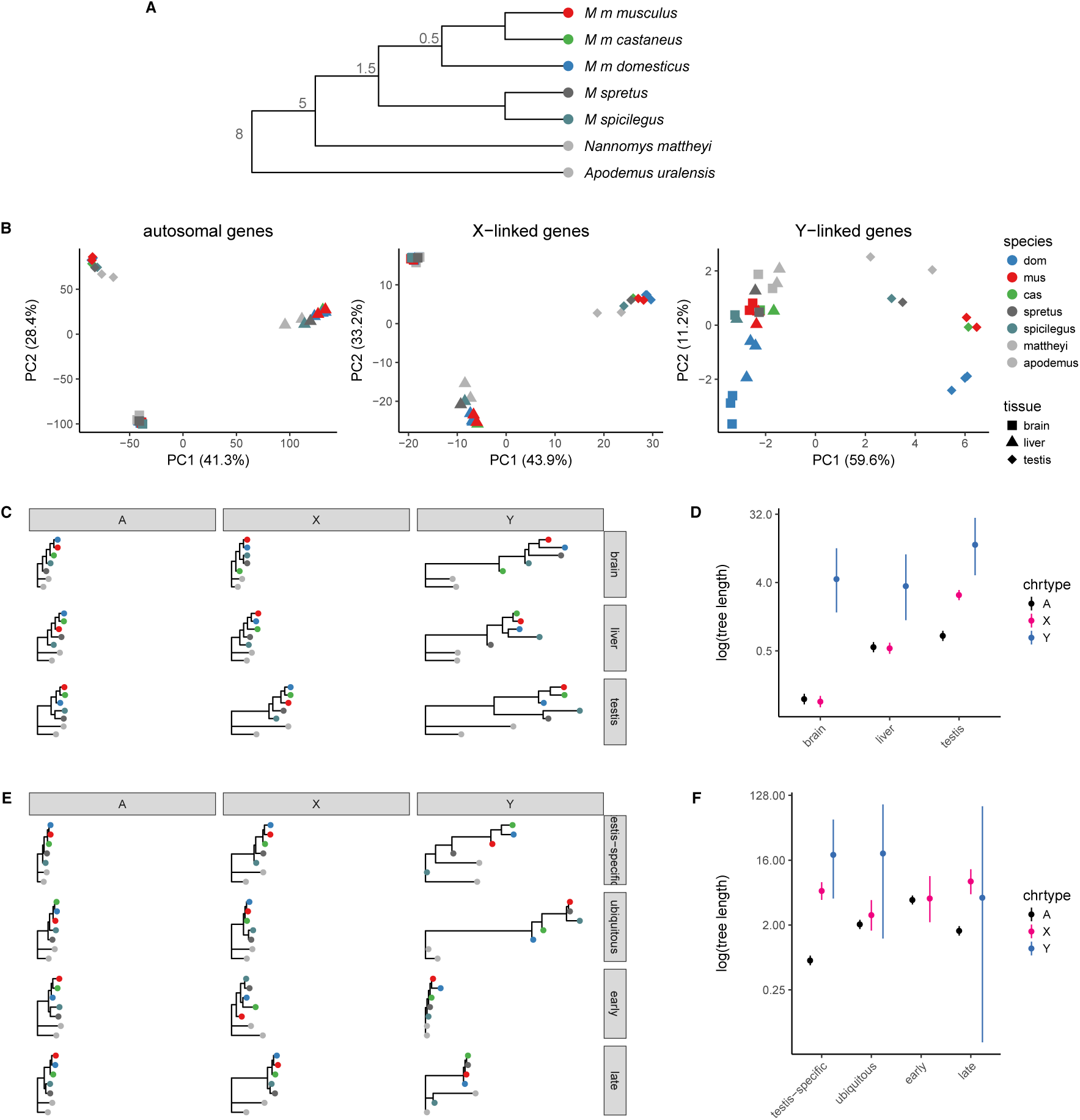
Divergence of sex-linked gene expression in murid rodents. (**A**) Schematic phylogeny of taxa in the multi-tissue expression dataset. Node labels are approximate divergence times (Mya); branch lengths not to scale. (**B**) Projection of samples onto the top two principal components of expression values for autosomal, X-linked and Y-linked genes. (**C**) Expression trees computed from rank-correlations between taxa for autosomal (A), X-linked (X) and Y-linked (Y) genes (across columns) for brain, liver and testis (across rows.) (**D**) Total tree length by chromosome type and tissue. (**E**) Expression trees as in panel C, with genes partitioned according to expression pattern: testis-specific; ubiquitously-expressed; early spermatogenesis (meiosis prior to MSCI); and late spermatogenesis (spermatids). (**F**) Total tree length by chromosome type and expression pattern.

To quantify divergence in gene expression patterns we computed the rank-correlation (Spearman’s *ρ*) between species for each tissue type separately for autosomal, X-linked and Y-linked genes, and constructed trees by neighbor-joining (Figure 9C). We use total tree length as an estimator of expression divergence. The topology of these trees for the autosomes and X chromosome in brain and testis is consistent with known phylogenetic relationships within the Muridae. Consistent with previous comparative analyses of gene expression in mammals (Brawand et al. 2011), we find that expression patterns are most constrained in brain and least constrained in testis (Figure 9D). Expression divergence is equal between autosomes and X chromosome in brain and liver, but greater for X-linked genes in testis. Y-linked expression diverges much more rapidly in all three tissues, but the effect is most extreme in the testis. We caution that the precision of these estimates is limited by the small number of Y-linked relative to autosomal or X-linked genes.

This “faster-X” effect should be limited to functional elements subject to male-specific selection. Genes expressed in the male germline (testis-biased and/or expressed during spermatogenesis) might be enriched for such elements, relative to genes with ubiquitous expression. We therefore estimated expression divergence in autosomal, X- and Y-linked genes with four sets of genes with different expression patterns (Figure 9E). X-linked expression diverges more rapidly than autosomal expression only among genes with testis-biased expression. In contrast to Larson et al. (2016b) but in keeping with other predictions (Good and Nachman 2005), we find that the “faster-X” effect on expression is larger for genes expressed late than early in meiosis (Figure 9F). The number of Y-linked genes in each group is too small to permit any strong conclusions.

## Discussion

We have shown that nucleotide diversity in *M. musculus* is reduced on both sex chromosomes relative to expectations for a stationary population, and that the effect appears strongest in *M. m. domesticus* and weakest in *M. m. musculus* (Table 3). Sex differences in the long-term average mutation rate, estimated from synonymous-sites divergence to rat, are not sufficient to explain the deficit. Because sex chromosomes respond differently than autosomes to changes in population size, we fit several (simple) models of demographic history to autosomal site-frequency spectra (Figure 7) and compared their predictions to observed values. At least for the models we considered (see **Supplement**), neither gradual nor instantaneous changes in population size — of magnitude feasible given autosomal SFS — can account for the reduction in diversity on both the X and Y chromosomes, even if we allow for a sex ratio different than 1:1 (Figure S3). Estimates of effective size of each population (from autosomal sites) are in agreement with previous work on house mice (Din et al. 1996; Baines and Harr 2007; Salcedo et al. 2007; Geraldes et al. 2008).

Using demographic histories from autosomes as a baseline, we simulated two modes of selection — background selection and hard selective sweeps — on Y chromosomes of *domesticus* and *musculus*. Although discrimination between models was limited by both technical factors and theoretical constraints, we have shown that the Y-linked SFS in *domesticus* is consistent with a moderately strong selective sweep (Figure S4). The background selection model is the best-fitting in *musculus*, but is only 1.4-fold more likely (log_10_ BF = 0.16) than the next-best model. We conclude that recent positive selection accounts, at least in part, for the reduction in Y-linked relative to autosomal diversity in *domesticus*. Furthermore, coding sequences of X-linked genes with germline expression are disproportionately shaped by positive selection (Figure 8). Both X- and Y-linked genes have rapidly-diverging expression patterns in the testis, especially in spermatids (Figure 9). Together these findings provide strong support for the idea that positive selection in the male germline is a potent and ongoing force shaping both mammalian sex chromosomes (Mueller et al. 2013; Larson et al. 2016b).

To what extent are these pressures a consequence of intragenomic conflict? The reciprocal actions of SLX/SLX1 and SLY on sex-linked gene expression in spermatids establish the conditions for conflict between the X and Y chromosomes that implicates *any* gene whose expression after meiosis is beneficial for sperm maturation and fertilizing ability. X-linked alleles that meet the functional requirement for post-meiotic expression in the face of repression by SLY — via a stronger promoter, a more stable transcript, a more active protein product, or increased copy number — should be favored by selection (Ellis et al. 2011). The same should be true, in reverse, for successful Y chromosomes.

Although we cannot directly identify the putative target(s) or meccanism(s) of selective sweeps on the Y chromosome, several independent lines of evidence point to the ampliconic genes on Yq active in the X-Y conflict. First, the copy number of *Slx/Slxl1* and *Sly* have increased three-fold within *M. musculus* and are correlated across populations (Figure 4), consistent with an “arms race” between the sex chromosomes in which the Y chromosome is the lagging player. The absolute expression of ampliconic X genes and their Yq homologs (in whole testis) increases with copy number across *Mus* (Figure S5). Larson et al. (2016a) have shown that, in spermatids from reciprocal *F*_1_ hybrids between *domesticus* and *musculus* that are “mismatched” for *Slx/Slxl1* and *Sly*, global X-chromosome expression is indeed perturbed in the direction predicted by the copy number and actions of SLX/SLX1 and SLY. Second, several independent deletions of Yq in laboratory stocks converge on a similar phenotype, namely low fertility, abnormal sperm morphology due to problems with chromatin compaction, and sex-ratio distortion in favor of females (Styrna et al. 1991; Conway et al. 1994; Touré et al. 2004; Fischer et al. 2016; MacBride et al. 2017). Third, Y chromosomes from *musculus* — the subspecies with highest *Sly* copy number — are more successful at introgressing across *domesticus-musculus* hybrid zone in Europe, and in localities where they do, the census sex ratio is shifted towards males (Macholán et al. 2008). Consomic strains with differing only by their Y chromosomes show similar deviation in the sex ratio from parity (Case et al. 2015). Finally, although modeling predicts moderately strong positive selection on Y, there is little evidence that it occurs within coding sequences of single-copy genes on Yp (Figure 8). This observation permits several explanations but is consistent with the idea that Yp alleles are hitchhiking with favorable alleles on Yq.

It is more difficult to ascertain the contribution of intragenomic conflict to the paucity of diversity on the X chromosome. Although the mammalian X chromosome is enriched for genes with expression in the male germline (*eg.* Rice (1984); Mueller et al. (2013)), its functional portfolio is considerably more broad than that of the Y chromosome (Bellott et al. 2014, 2017). The X chromosome also has a major role in hybrid sterility in mouse (Forejt and Iványi 1974; Forejt 1996; Storchová et al. 2004; Payseur et al. 2004; Teeter et al. 2008; Good et al. 2008; Campbell et al. 2013; Turner et al. 2014); the Y chromosome does not (Turner et al. 2012; Campbell and Nachman 2014). We corroborate the “faster-X” effect on protein evolution that has been previously described by others (Kousathanas et al. 2014; Larson et al. 2016b) and show that it is strongest for genes expressed in the male germline (Figure 8), which are widely scattered across the X chromosome (Figure S2D). We conclude that selection is pervasive on the mouse X chromosome and reduces diversity chromosome-wide. This stands in contrast to the pattern observed in great apes, which has apparently been driven by a few strong selective sweeps (Hvilsom et al. 2012; Veeramah et al. 2014; Nam et al. 2015).

Many open questions remain with respect to the evolution of mouse Y chromosomes. How many of the hundreds of copies in each gene family retain coding potential? Which copies are functionally equivalent? Does suppression of recombination promote the spread of clusters of genes like *Slx*, similar to sex-ratio drivers in other species (Jaenike 2001)? What evolutionary trade-offs does success in the sex-chromosome conflict entail, in the context of sperm competition and polyandry in natural populations (Simmons and Fitzpatrick 2012)? Does the conflict lead to oscillations between a male-biased and female-biased population over time, and if so, what is the effect on patterns of diversity on the sex chromosomes? All of these are important avenues of future study as we seek to understand the forces shaping sex chromosomes.

## Materials and methods

### Alignment and variant-calling

Whole-genome sequencing reads were obtained from the European Nucleotide Archive (PRJEB9450, PRJEB11742, PRJEB14673, PRJEB14167, PRJEB2176, PRJEB15190) and whole-exome reads from the NCBI Short Read Archive (PRJNA323493). Reads were aligned to the mm10 reference sequence using bwa mem v0.7.15-r1140 (Li 2013) with default parameters. Optical duplicates were marked using samblaster and excluded from downstream analyses. Regions of the Y chromosome accessible for variant calling were identified using the CallableLoci tool in the GATK v3.3-0-g37228af (McKenna et al. 2010). To be declared “callable” within a single sample, sites were required to have depth consistent with a single haploid copy (3 < depth < 50) and < 25% of overlapping reads having mapping quality (MQ) zero. The analysis was restricted to Yp. The final set of callable sites was defined as any site counted as callable within > 10 samples. In total, 2 289 336 bp (77% of the non-gap length of Yp) were deemed callable.

SNVs and short indels on the Y chromosome were ascertained using GATK HaplotypeCaller v3.3-0-g37228af in the intersection of callable regions and exons targeted by the Roche NimbleGen exome-capture array, lifted over to mm10 with CrossMap v0.2.3 and the mm9-to-mm10 chain file from the UCSC Genome Browser (http://hgdownload.soe.ucsc.edu/goldenPath/mm9/liftOver/mm9ToMm10.over.chain.gz). To minimize artifacts from cryptic copy-number variation, X-Y homology, and the like, only biallelic sites with a “homozygous” (*ie.* single-copy hemizygous) call in all male samples were used. Sites called in fewer than 60 samples or with strand-bias *p*-value < 0.01 were filtered. Raw VCF files are provided in **File S1**.

For the Y chromosome phylogenetic tree shown in Figure 2A, data from Collaborative Cross lines carrying A/J, 129S1/SvImJ, NOD/ShiLtJ, NZO/HlLtJ, CAST/EiJ, PWK/PhJ, and WSB/EiJ Y chromosomes were used in place of the inbred strains themselves. (See aliases in **Table S1**.) Whole-genome sequence from male representatives of these lines has not (to our knowledge) been published.

### Estimation of site frequency spectra and summary statistics

Site frequency spectra (SFS) were calculated from genotype likelihoods at callable sites using ANGSD v0.917 (Korneliussen et al. 2014). Genotype likelihoods for the autosomes were caculated under the GATK diploid model after applying base alignment quality (BAQ) recalibration with the recommended settings for bwa alignments (-baq 1 -c 50, effectively discarding evidence from reads aligning at < 95% identity). Sites were filtered to have per-individual coverage consistent with the presence of a single diploid copy (3 < depth < 80), to be non-missing in at least 3 individuals per population. Genotype likelihoods for the X and Y chromosomes were calculated under the GATK haploid model with depth filters appropriate for haploid sites (3 < depth < 40). Only reads with MQ > 20 and bases with call quality > 13 were considered, and ampliconic regions (plus a 100 kb buffer on each side) were masked. Site-wise allele frequencies were computed within each population separately, and the joint SFS across non-missing sites in the three populations was estimated from these frequencies. The consensus genotype from a single *Mus spicilegus* male was used as the ancestral sequence to polarize alleles as ancestral or derived. Ensembl v87 reference annotations were used to define internecine sites, intronic sites, 0-fold and 4-fold degenerate sites.

Diversity statistics and neutrality tests were calculated from joint SFS using standard formulae implemented in a custom Python package, sfspy (http://github.com/andrewparkermorgan/sfspy). Uncertainties for autosomal and X-linked sites were obtained by bootstrapping over loci, since the X and autosomes recombine; and for Y-linked sites using the built-in bootstrapping method of ANGSD.

### Models of sex-chromosome diversity under neutral coalescent

The expected ratio of X-to-autosome (X:A) and Y-to-autosome (Y:A) pairwise diversity was obtained from the formulae derived in Pool and Nielsen (2007). Define the inheritance factors *h*_*A*_ = 1, *h*_*X*_ = 3/4 and *h*_*Y*_ = 1/4; and mutation rates *μ*_*A*_, *μ*_*A*_, *μ*_*Y*_. For an instantaneous change in population size of depth *f* from starting size *N*, the expected value of X:A is:

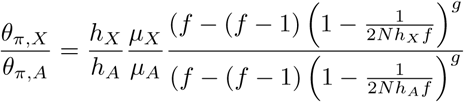

The expression for Y:A can be written similarly. Note that X:A and Y:A depend only on the ratio between mutation rates on different chromosomes, not the absolute mutation rate. For a bottleneck of depth *f*, starting *g*,_1_ generations before the present and ending at *g*_1_ + *g*_2_ generations before the present:

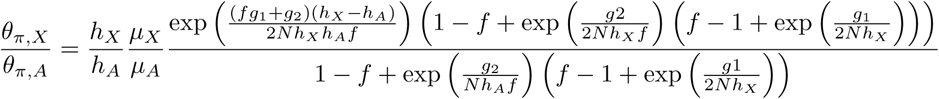

For a model with exponential growth with rate constant *r*, we used the approximation provided in Polanski et al. (2017):

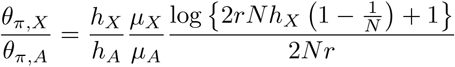

Unequal sex ratios were modeled by calculating the number of X and Y chromosomes per autosome, given fixed autosomal effective population size, using standard formulae as in Sayres et al. (2014), and passing these into the equations above via the parameter *h*_*X*_ or *h*_*Y*_.

In Figure S3, we plot X:A against Y:A. For the bottleneck and step-change models, X:A and Y:A vary with time since the onset of size change; these trajectories can be traced clockwise along each curve from *t* = 0 to *t* = 4*N* (backwards in time.)

### Demographic inference

The four demographic models illustrated in Figure 7A were fit to autosomal SFS using *∂a∂i* v1.7 (Gutenkunst et al. 2009). We fit each model separately to each population, using the sum of marginal spectra from 1000 approximately unlinked, putatively neutral intergenic regions each 100 kb in size, spanning a total of 85.4 Mb of callable sites after removing those missing in one or more populations. Because the depth, duration and onset of a bottleneck have are confounded in the SFS, we fixed the duration of the bottleneck to be short (0.1*N*_*e*_ generations) and attempted to estimate the remaining two parameters. We additionally constrained the bottleneck model to include recovery to exactly the starting population size.

Convergence of model fits was assessed qualitatively by re-fitting each model from 10 sets of randomly-drawn initial values. We confirmed that the best-fitting models shown in Figure 7 represent the “modal” result, in that a majority of independent runs reach a solution within 5 log-likelihood units of the one shown. Parameter estimates should nonetheless be interpreted with caution, as their uncertainties are wide.

### Models of natural selection

To model the effect of natural selection on Y-linked diversity while accounting for possible non-stationary demographic processes, we used forward simulations implemented in SLiM v2.2.1 (Haller and Messer 2017). For *M. m. domesticus* we simulated from a stationary model; for *M. m. musculus*, from an exponential growth model.

We considered two modes of selection: background selection (BGS) due to purifying selection against deleterious mutations at linked sites; and hard selective sweeps on newly-arising beneficial mutations. Relative fitness in SLiM is modeled as 1 + *s* for sex-limited chromosomes. BGS was modeled by introducing mutations whose selection coefficients *s* were drawn from a mixture of a gamma distribution with mean *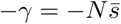* (100 *× α*% of mutations), and a point mass at zero ((100 × (1 − *α*)% of mutations.) BGS simulations were run for 10*N* generations, sufficient to reach mutation-selection-drift equilibrium. For selective sweeps, the simulation was first run for 10*N* generations of burn-in, and then a beneficial variant was introduced with *s* drawn from a gamma distribution with mean *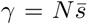*. The simulation was then tracked until the beneficial variant was fixed or lost; in the case of loss, the run was re-started from the end of the burn-in period with a new mutation. We confirmed the integrity of simulations by checking that the pairwise diversity achieved by runs with selection coefficients fixed at zero matched the observed neutral values for each population (not shown.) Values of *α* were drawn from a uniform distribution on (0, 1], and values of *γ* were drawn from a log-uniform distribution on (10^−6^, 103]. Runs were scaled for computational performance.

Simulations were connected to an approximate Bayesian computation (ABC) inference procedure implemented with the R package abc (Csilléry et al. 2012). Briefly, 500, 000 simulations were performed for each model. Five summary statistics were calculated from the SFS generated by each simulation: Watterson’s estimator *θ*_*w*_; Tajima’s estimators *θ*_*π*_ and *θ*_*ζ*_; Tajima’s *D*; and Fu and Li’s *D*. The same set of statistics was computed for the observed joint SFS. The 0.1% of simulations with smallest Euclidean distance to the observed summary statistics were retained, accounting for collinearity between summary statistics using the “neuralnet” method of the function abc::abc(). Posterior distributions were computed via kernel smoothing over the parameter values of the retained simulations using an Epanechnikov kernel and plug-in bandwidth estimate.

Models were compared via their Bayes factors, calculated using the abc::postpr() function. To confirm the fidelity of the best-fitting model, summary statistics for pseudo-observed datasets (*i.e.* simulations from the posterior distributions) were checked against the observed summary statistics.

### Size estimation of co-amplified regions of Yq and X

Copy number of ampilconic genes on Yq and X was estimated as follows. First, all paralogs in each family were identified by BLAT and BLAST searches using the sequences of canonical family members from Ensembl. These searches were necessary because many member of each family are annotated only as “predicted genes” (gene symbols “GmXXXX”). Based on BLAST results we assigned the *Spin2/4* family – with members in several clusters on the proximal X chromosome – as *Sstx*. Normalized coverage was estimated for each non-overlapping paralog by counting the total number of reads mapped and dividing by the genome-wide average read depth.

### Identification of *de novo* CNVs in Collaborative Cross lines

Whole-genome sequencing reads (2 × 150 bp paired-end) from a single male individual from each 69 distinct Collaborative Cross (CC) lines were obtained from Srivastava et al. (2017). Alignment and quality control was performed as for wild mice. Read depth was estimated in 100 kb bins across the Y chromosome for each individual, and normalized for the effective depth of sequencing in that sample. Unambiguous alignment of 150 bp reads to the highly repetitive sequence on Yq is clearly not possible. However, each of the 8 founder Y chromosome haplogroups in the CC produces a characteristic read depth profile when reads are aligned with bwa-mem. We exploited this fact to remove noise from ambiguous read mapping by re-normalizing the estimated depth in each bin for each sample agains the median depth in that bin for CC lines sharing the same Y chromosome haplogroup (listed in **Table S2**). Any remaining deviations in read depth represent variation among lines sharing the same Y chromosome haplogroup, that is, candidate *de novo* CNVs. CNVs were ascertained by manual inspection of the re-normalized read depth profile of each CC line.

### Analyses of gene expression

*Multi-tissue, multi-species dataset.* Neme and Tautz (Neme and Tautz 2016) measured gene expression in whole testis from wild-derived outbred mice from several species (Figure 9A) using RNA-seq. Reads were retrieved from the European Nucleotide Archive (PRJEB11513). Transcript-level expression was estimated using kallisto (Bray et al. 2016) using the Ensembl 85 transcript catalog augmented with all *Slx/y*, *Sstx/y* and *Srsx/y* transcripts identified in (Soh et al. 2014). In the presence of redundant transcripts (*i.e.* from multiple copies of a coamplified gene family), kallisto uses an expectation-maximization algorithm to distribute the “weight” of each read across transcripts without double-counting. Transcript-level expression estimates were aggregated to the gene level for differential expression testing using the R package tximport. As for the microarray data, “predicted” genes (with symbols “GmXXXX”) on the Y chromosome were assigned to a co-amplified family where possible using Ensembl Biomart.

Gene-level expression estimates were transformed to log scale and gene-wise dispersion parameters estimated using the voom() function in the R package limma. Genes with total normalized abundance (length-scaled transcripts per million, TPM) < 10 in aggregate across all samples were excluded, as were genes with TPM > 1 in fewer than three samples.

*Spermatogenesis time course.* Larson et al. (2016a) measured gene expression in isolated spermatids of three males from each of four *F*_1_ crosses — CZECHII/EiJ×PWK/PhJ; LEWES/EiJ×PWK/PhJ; PWK/PhJ×LEWES/EiJ; and WSB/EiJ×LEWES/EiJ — using RNA-seq. Reads were retrieved from NCBI Short Read Archive (SRP065082). Transcript-level expression was estimated using kallisto (Bray et al. 2016) using the Ensembl 85 transcript catatlog augmented with all *Slx/y*, *Sstx/y* and *Srsx/y* transcripts identified in (Soh et al. 2014). In the presence of redundant transcripts (*i.e.* from multiple copies of a co-amplified gene family), kallisto uses an expectation-maximization algorithm to distribute the “weight” of each read across transcripts without double-counting. Transcript-level expression estimates were aggregated to the gene level for differential expression testing using the R package tximport. As for the microarray data, “predicted” genes (with symbols “GmXXXX”) on the Y chromosome were assigned to a co-amplified family where possible using Ensembl Biomart.

Gene-level expression estimates were transformed to log scale and gene-wise dispersion parameters estimated using the voom() function in the R package limma. Genes with total normalized abundance (length-scaled transcripts per million, TPM) < 10 in aggregate across all samples were excluded, as were genes with TPM > 1 in fewer than three samples.

*Definition of tissue-specific gene sets.* The “tissue specificity index” (*τ*) of Yanai et al. (2005) was used to define tissue- or cell-type-specific gene sets. The index was first proposed for microarray data, and was adapted for RNA-seq as follows. Define *T*_*i*_ to be the mean log-scaled expression of a gene in tissue or cell type *i* (of *N* total), as estimated by limma. We require expression values to be strictly positive, so let *q* = min_*i*_ *T*_*i*_ and define *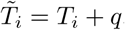*. Finally, calculate *τ* as

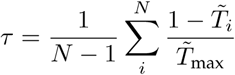

The set of testis-biased genes was defined as all those with *τ* > 0.5 and higher expression in testis than in any of the other seventeen tissues in the multi-tissue dataset (PRJEB11897, Harr et al. (2016)). The set of ubiquitously-expressed genes was defined as those with *τ* < 0.25 and whose expression was above the median expression in the highest-expressing tissue. The set of early-meiosis genes was defined as those with *τ* > 0.5 and highest expression in leptotene/zygotene spermatocytes; spermatid genes were defined as those with *τ* > 0.5 and highest expression in round spermatids. We analyzed expression specificity during spermatogenesis separately in the two intra-subspecific *F*_1_ crosses, and took the union of the resulting gene sets.

## Acknowledgments

The authors thank Jeff Good, Erica Larson, Michael Nachman, Megan Phifer-Rixey, Jacob Mueller, Alyssa Kruger, Marty Ferris, Peter Ellis and additional anonymous reviewers for many insightful comments and suggestions. This work was supported by the National Institutes of Health (F30MH103925, R01HD065024, U19AI100625, U42OD010924).

## Supplement

### Caveats to demographic models

Like all models, the scenarios of neutral demography and natural selection presented here are greatly simplified and almost certainly wrong. We chose to consider the history of each of the three subspecies independently, rather than in a joint isolation-with-migration or isolation-by-distance model, because the one-population models could be fit more robustly and more easily checked against analytical formulae. The simple models are nonetheless useful as guideposts along the way to a better approximation of the true history of the mouse sex chromosomes. On the basis of autosomal SFS, we can confidently reject scenarios with a single very sharp (*f* ≪ 0.1) reduction in population size with or without unequal sex ratio as the sole explanation for lack of Y-linked diversity. Likewise we can rule out exponential growth alone, because it actually increases X:A and Y:A relative to a stationary model (Figure S3C). One important possibility that we have not considered is a fluctuating sex ratio. An “arms race” between the X and Y chromosomes for transmission in the male germline could lead to oscillations between a male-biased and female-biased population. Even if deviations in the sex ratio are transient, the net effect could be to reduce diversity on both sex chromosomes relative to autosomes out of proportion to the strength of selection.

Disentangling the effects of demography and selection on the Y chromosome is especially challenging because the Y has the smallest effective population size and is inherited without recombination, so it is most susceptible to changes in population size, to background selection and to selective sweeps. We have used ABC to show that Y-linked SFS are consistent with recent positive selection. Of course the fact that selective sweeps offer plausible fit, conditional on neutral demographic history, does not rule out background selection. Both mammalian Y and avian W chromosomes, which have independent evolutionary origins, retain a convergent set of dosage-sensitive genes with roles in core cellular processes (Bellott et al. 2017). The proportion of substitutions in these genes fixed by positive selection (*α*_SEW_, Figure 8) is indeed much smaller than for X-linked genes. Together these results imply relatively strong purifying selection on ancestral Y genes, which in the absence of recombination should constrain diversity on the entire chromosome. This is an important alternative hypothesis to a hitchhiking effect associated with positive selection on Yq.

### Supplementary tables

**File S1.** Raw VCF files with genotype calls on Y chromosome and mitochondrial genome, available from the Zenodo repository: https://doi.org/10.5281/zenodo.817658.

**Table S1.** List of samples used in this study (Excel spreadsheet). Sample manifest provided in first tab, column key in second tab.

**Table S2.** Y chromosome haplogroup assignment and CNV status for 69 Collaborative Cross strains.

**Table S3.** Sequence diversity statistics across different classes of sites on the autosomes, X and Y chromosomes, by population. See main text for details. *L*, total number of callable bases in target locus; *θ*_*π*_, Tajima’s pairwise *θ*; *θ*_*w*_, Watterson’s *θ*; *D*, Tajima’s *D*; *D*_*F* *L*_, Fu and Li’s *D*. Both estimators of *θ* are expressed as percentages with bootstrap standard errors in parentheses.

### Supplementary figures

**Figure S1:**
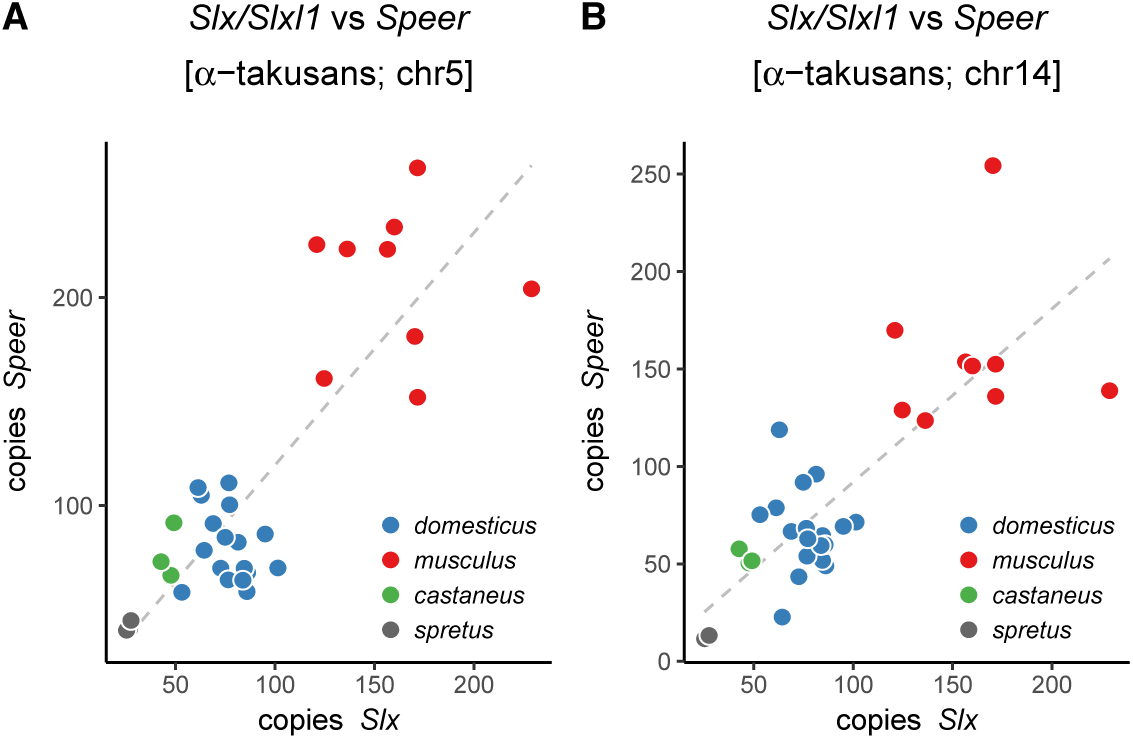
Copy number of *Speer* family members on chromosomes 5 (**A**) and 14 (**B**) compared to copy number of *Slx/Slxl1*.

**Figure S2:**
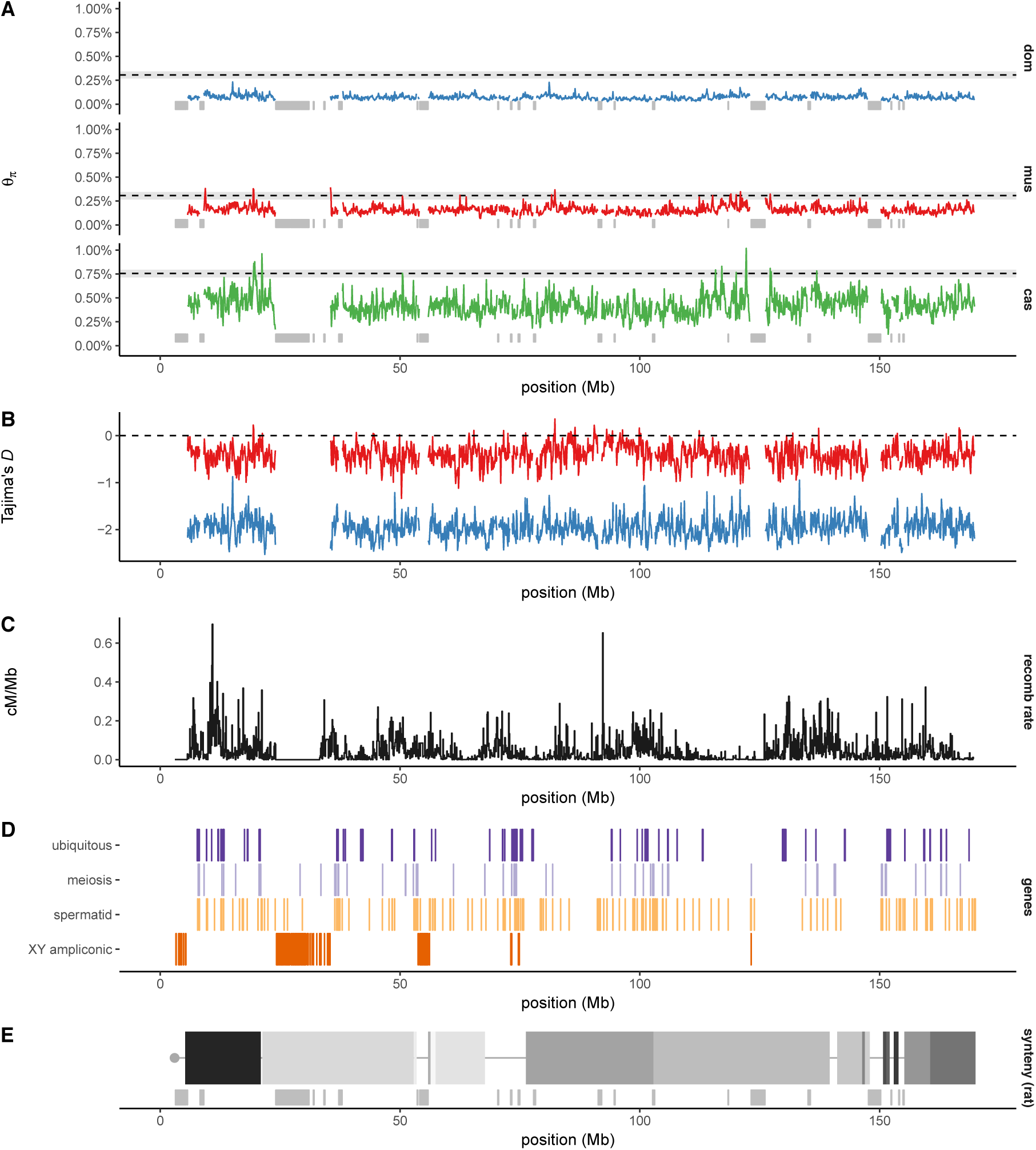
Global view of sequence diversity on the X chromosome. (**A**) Pairwise diversity in 100 kb windows by population, with expectation based on autosomal diversity shown as dashed line with 95% confidence band. Ampliconic regions from Mueller et al. (2008) were masked and are indicated with grey bars. (**B**) Tajima’s *D* for *domesticus* and *musculus*, with masking as in panel A. (**C**) Recombination rate (centimorgans per Mb) estimated in the Diversity Outbred stock. (**D**) Locations of gene sets used elsewhere in this manuscript, plus X-Y ampliconic genes from Soh et al. (2014), as sketched in Figure 1. (**E**) Blocks of conserved synteny with rat. Boundaries between blocks define rearrangement breakpoints between mouse and the common rodent ancestor.

**Figure S3:**
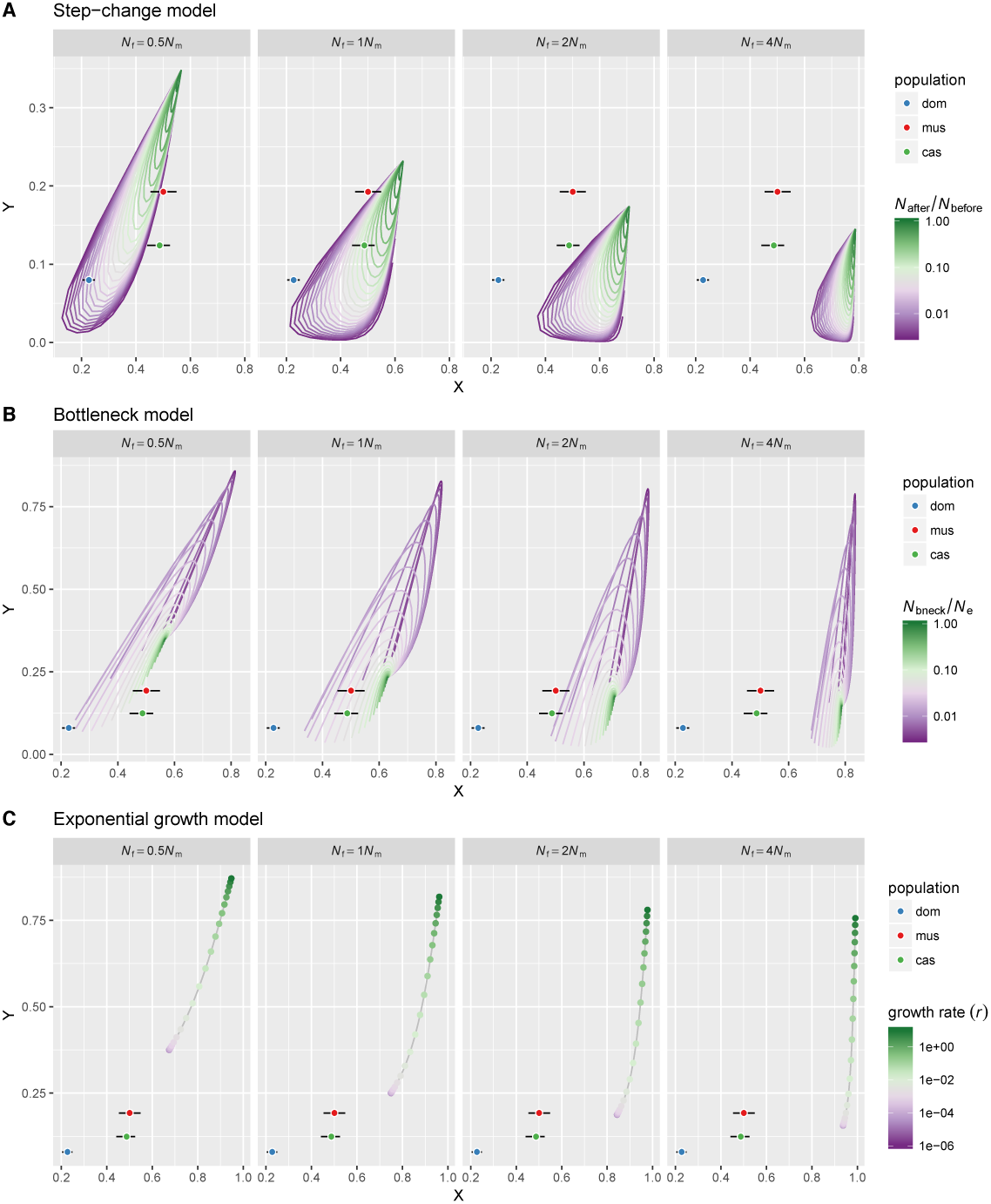
X:A and Y:A diversity ratios under several neutral coalescent models with varying sex ratio. Error bars represent bootstrap standard errors.

**Figure S4:**
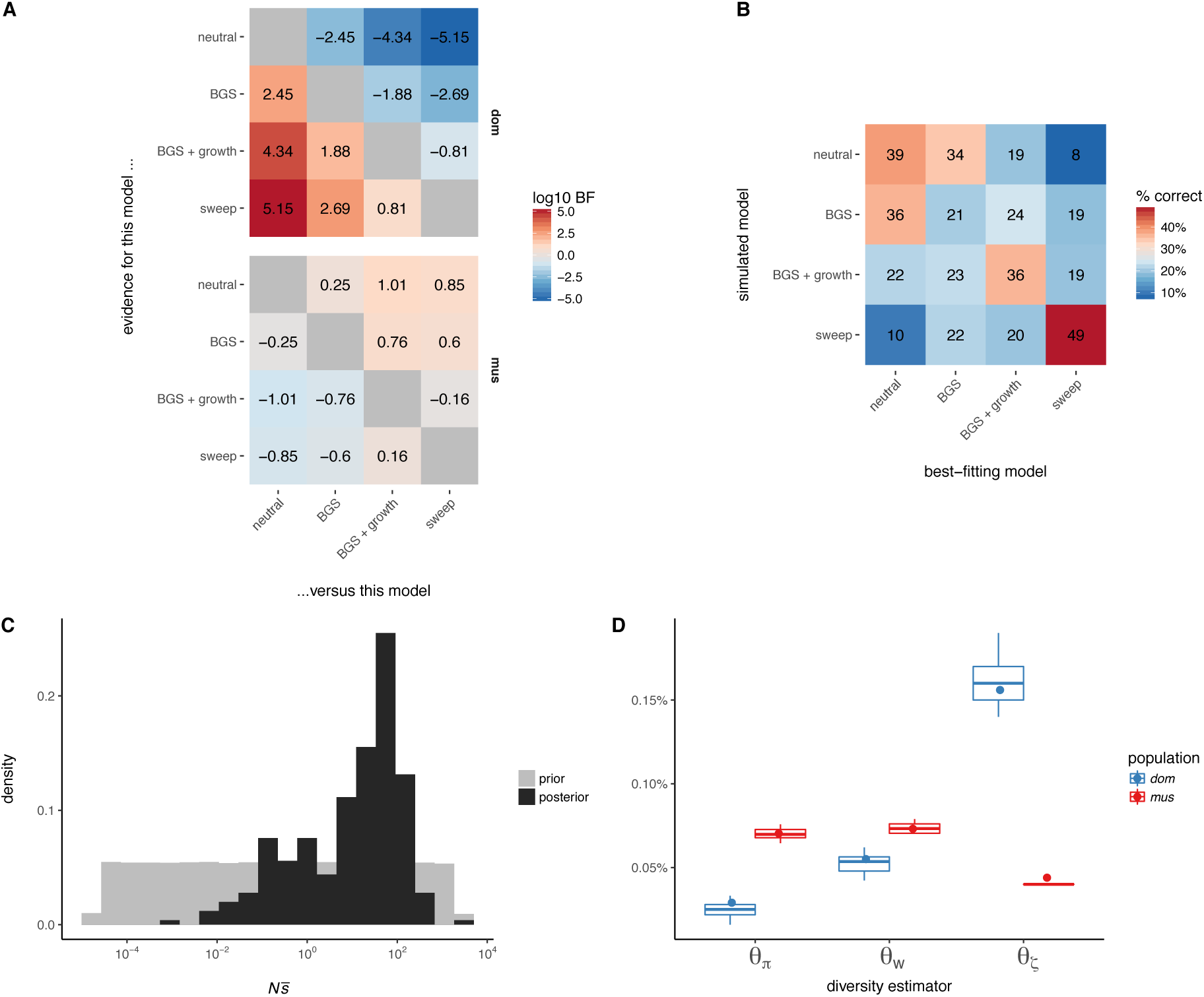
Modelling selection on the Y chromosome. (**A**) Pairwise comparison of goodness-of-fit using Bayes factors. (**B**) Percent recall of model-selection procedure estimated by leave-one-out cross-validation. (**C**) Prior and posterior distributions of the population-scaled mean selection coefficient 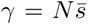 for the selective-sweep model in *domesticus*. (**D**) Posterior distribution of diversity statistics from the best-fitting model in each population (boxplots), compared to their observed values.

**Figure S5:**
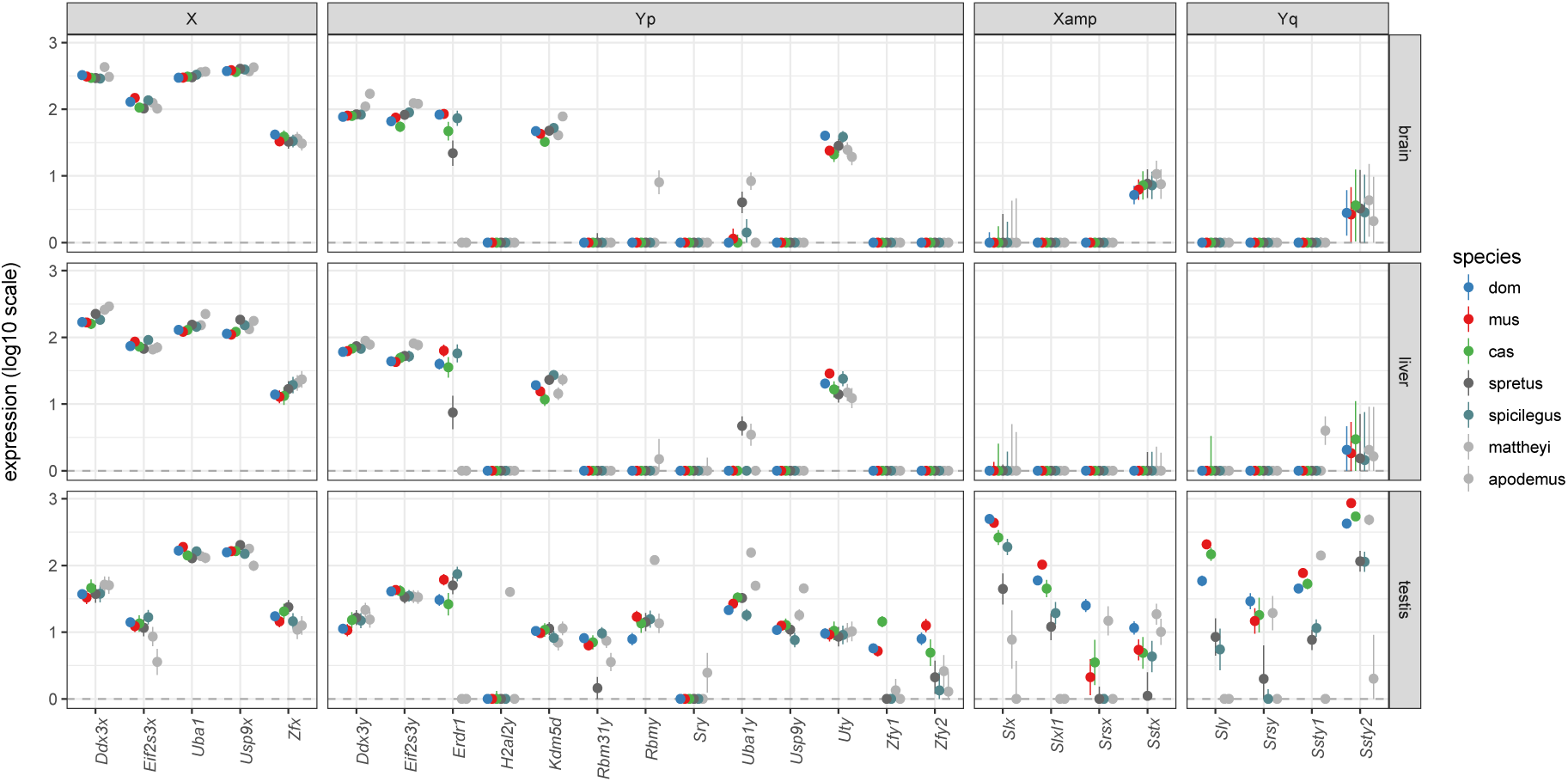
Expression of selected X- and Y-linked genes across *Mus* species. Genes are subdivided by chromosomal location: X, non-ampliconic X-linked genes with Y-linked homologs; Yp, non-ampliconic genes on Yp; Xamp, X-linked homologs of co-amplified genes; Yq, Y-linked homologs of co-amplified genes, residing on Yq.

